# Mutation rate heterogeneity at the sub-gene scale due to local DNA hypomethylation

**DOI:** 10.1101/2023.09.26.559585

**Authors:** David Mas-Ponte, Fran Supek

## Abstract

Local mutation rates are highly heterogeneous across the human genome. This variability was better studied at the scale of megabase-sized chromosomal domains on the one extreme, and at the scale of oligonucleotides at the other extreme. The intermediate, kilobase-scale heterogeneity in mutation risk was less studied. Here, by analyzing thousands of somatic genomes, we considered the hypothesis there are mutation risk gradients along gene bodies, representing a genomic scale spanning roughly 1 kb – 10 kb, and that different mutational mechanisms are differently distributed across gene segments. The main intragenic heterogeneity concerns several kilobases at the transcription start site and further into 5’ ends of gene bodies, which are commonly hypomutated with respect to several mutational signatures, most prominently the ubiquitous mutational signature of C>T changes at CpG dinucleotides. Width and shape of this mutational coldspot at 5’ gene ends is variable across genes, and corresponds to variable interval of lowered DNA methylation across genes. These hypomutated genic intervals correspond to hypomethylation that can originate from various causes, including intragenic enhancers, Polycomb-marked regions, or chromatin loop anchor points. Tissue-specific DNA hypomethylation begets tissue-specific local hypomutation. However, direction of mutation rate effect is inverted for some mutational processes, where signatures of AID/APOBEC3 cytosine deaminase activity are actually increased in hypomethylated regions. Overall, local DNA methylation determines mutation rate heterogeneity at the sub-gene level, and can generate either mutational coldspots or hotspots, depending on the mutagen exposure history of a cell.

## Introduction

The local variation in mutation rates along the human genome is evident at different scales^1^. At the coarse resolution, mutation rates vary substantially across approximately megabase-sized domains^2,3^, which correspond to replication timing domains and to topologically-associated domains^4,5^. This heterogeneity is generated by the differential activity of DNA repair pathways^6,7^, which lower mutation rates in early-replicating, euchromatic domains. At the fine resolution, mutation rates vary strongly according to the trinucleotide sequence, yielding patterns of sequence predisposition termed mutation signatures^8–10;^ moreover the pentanucleotide and heptanucleotide sequence neighborhoods predict mutation risk for some processes^11,12^. In addition, individual examples of other factors that have an intermediate resolution in the genome can modify the accumulation of mutations by interacting either directly or indirectly with DNA damage and repair processes^1^. This includes for instance nucleosome positioning^13^, secondary DNA structures^14,15^, CTCF^16–18^ and ETS transcription factor binding^19–22^, and locally open, accessible chromatin^23–25^. These individual examples suggest there may be additional, extensive heterogeneity in mutation rates below the domain-scale and above the oligonucleotide scale (the range may be referred to as “mesoscale”^14,26^).

We were motivated to consider mutation rate gradients across gene bodies, because some epigenetic features, such as histone modifications^2^ are known to be variably deposited along genes and also known to associate with mutation rates. For instance, H3K36me3 mark is deposited variably along the length the transcribed gene bodies^27^. This mark recruits the DNMT3B^28,29^ protein, which increases gene body DNA methylation, and also the DNA repair protein MSH6^30^; both factors have potential to affect mutation rates^31^. The H3K79 methylation mark was also reported to accumulate differentially across gene bodies^32^ and may participate in the repair of breaks, UV damage and the control of error-prone DNA polymerases^33–35^, thus with potential to affect mutation rates.

In addition to histone modifications, DNA methylation at the cytosine in CpG dinucleotides^36^ is pervasive in the human genome, and is mutagenic since spontaneous deamination directly generates a T/G mismatch^37^. A mismatch during DNA replication of the methylated cytosine has also been proposed as a mechanism of mutagenesis^38^, particularly upon DNA repair failures^39–41^. CpG dinucleotides are variably distributed across the genome, locally accumulating at CpG islands near gene transcription start sites (TSS), which regulate gene transcription via hypomethylation or hypermethylation of promoter-proximal DNA^36^. Thus, the enrichment of the highly mutable CpG context near transcription start sites, as well as its variable DNA methylation, also represents a potential local mutation risk modifier at the gene scale.

Here, we perform a systematic and unbiased analysis of the mutation rate gradients along gene bodies, highlighting a major role of DNA methylation in generating genome-wide heterogeneity in local mutation risk, which differs across mutational signatures. We also quantify the role of DNA hypomethylation in other functional elements, such as enhancers and chromatin loop anchors, which also exhibit hypomethylation and constitute coldspots of certain mutational processes also when not associated with genes, further underscoring the role of DNA methylation in generating local mutation rate variation in the human genome.

## Results

### Sub-gene mutation rate gradients are observed in DNA methylation-associated mutational signatures

To systematically analyze the sub-gene variability of mutation rate within genes, we used a set of 2,782 tumor and healthy somatic whole-genome sequences (see Methods and Supp. Table 1), mainly from the PCAWG dataset^42^ and other publicly available sources^43–45^, to calculate the relative mutation enrichment at various loci across the length of gene bodies. In brief, we separated mutations by the mutational signature likely to have generated them (see Online Methods) and binned the gene body into 250bp long segments covering an extended (6kb long) region along both the 5’ and 3’ gene ends and additionally a central region (2kb long). Genes were pooled for this analysis and were considered stratified into three bins by their average mRNA expression levels across various tumor types (see Online Methods). We estimated the local relative mutation rate, controlling for variable trinucleotide composition of each 250bp long segment, using a negative binomial regression approach (see Online Methods and Fig. 1A).

**Fig. 1.**
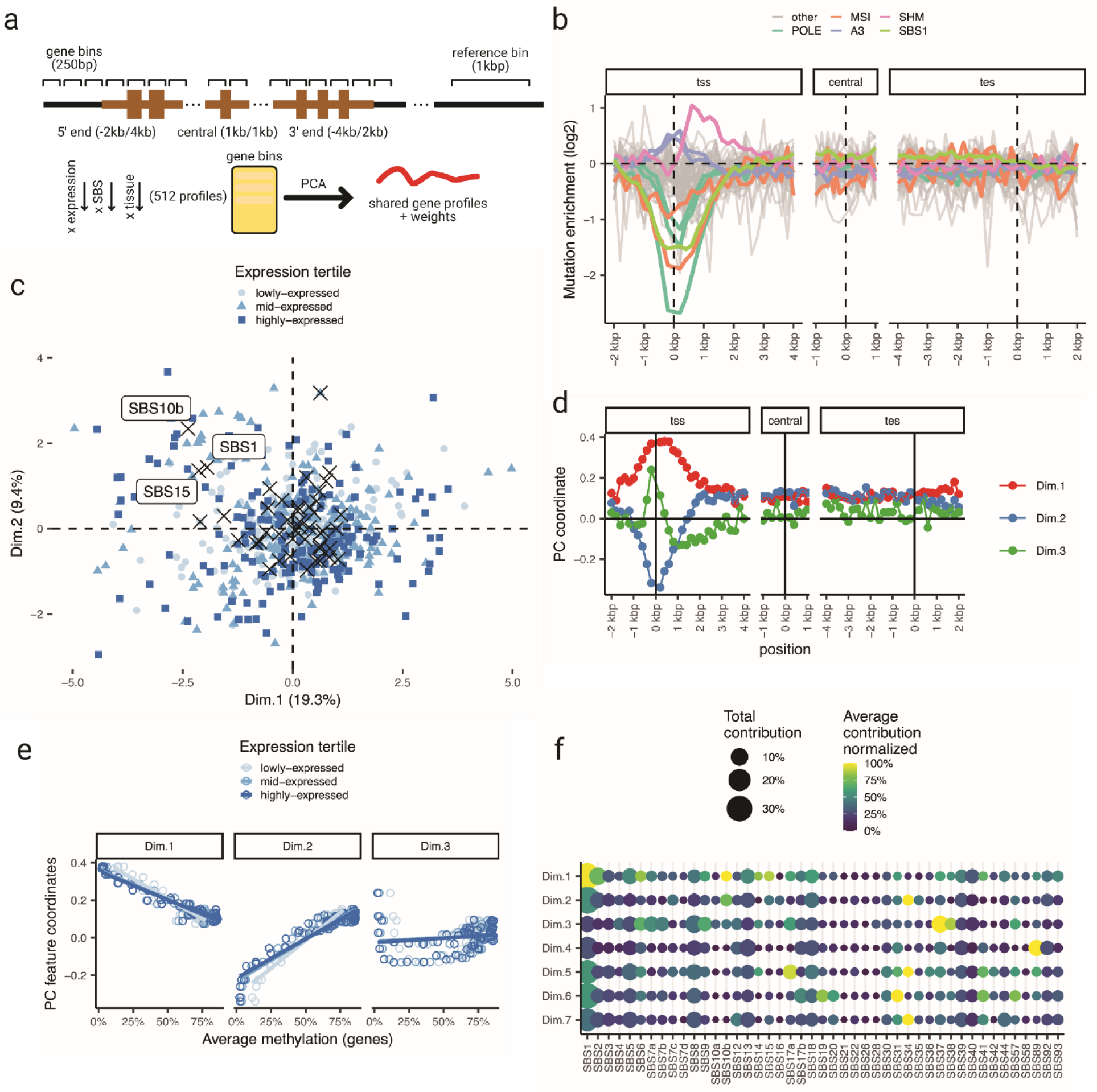
Systematic quantification of mutation gradients along gene bodies. (a) Diagram of the analysis of mutation rate gradients process and posterior factorization with a PCA. The genes are divided in 250bp long bins for which the mutation rate is calculated. The mutation rates at each bin is measured with a negative binomial regression and the output is factorized using a PCA. (b) Mutation rate estimates for different mutational signatures in a pan-cancer setting. Gene regions defined as 250bp bins across extended regions near both genic ends and the central position. (c) PCA coordinates of the instances included in the regression, here 512 points representing each combination of expression bin, signature and tissue of origin. (d) Profile weights of the three principal component along the gene body. (e) Correlation of median methylation levels across genes against coordinates of component 1, 2 and 3. (f) Contribution of each mutational signature to each component. In color the average contribution within the tissues that contain a certain mutational signature is represented while the size represents the total contribution against all cancer types.

Upon visual inspection of the pan-cancer mutation rate profiles, stratified by signature (averaged over expression bins, Fig. 1B and Supp. Fig. 1) we observed a strong and consistent mutation rate depletion at the 5’ gene end, occurring mainly for some signatures: SBS1, associated with the deamination of methylated cytosines; SBS15 and SBS6, associated with the MMR deficiencies (microsatellite instability, MSI), and SBS10a, SBS10b and SBS28, associated with mutations in the proofreading domain of the replicative DNA polymerase ε (*POLE*)^46,47^. In contrast to this depletion, the APOBEC3-associated mutation signatures, SBS2 and SBS13, showed an increase in the region surrounding the transcription start site (TSS). As a positive control, we observed that SBS9, which associates with somatic hypermutation (SHM) process in lymphoma, presented a strong enrichment in the 5’ gene ends, consistent with known localization of SHM target loci near active promoters (Fig. 1B)^48^. However, when considering the central section of the gene or the 3’ gene ends, in most mutational signatures no clear trends were observed. Overall, in a pan-cancer setting, most of the heterogeneity in mutation rates at the sub-gene resolution results in gradients at the 5’ gene end.

To more rigorously study these trends and their association with tissue and with gene activity levels, we next extracted the dominant patterns in mutation enrichment along the selected (central, 5’ and 3’ ends) sub-regions of the gene bodies using a principal component (PC) analysis (Fig. 1A,C); here, the mutation data was further divided by tissue-of-origin to confirm that trends are consistent across cancer types (Supp. Fig. 2). The first PC accounted for ∼19% of the systematic variability in trinucleotide-adjusted local mutation rates (Extended Data Fig. 1A); its profile along the gene body plateaued at the TSS and the 1 kb downstream thereof, and continued in the downstream regions, with some signal apparent 3kb downstream of TSS (Fig. 1D). The second component (PC2), explaining ∼9% variability in local mutation risk (Extended Data Fig. 1A), showed a narrower peak at the TSS loci (note that the overall inverse direction *versus* PC1 is arbitrarily assigned by PCA; Fig. 1D). We interpret the first two PCs jointly: their combination describes the variation in the width and/or depth of the 5’ hypomutated region across genes and/or mutation signatures. This PC analysis confirms that most of the variability in mutation rates within genes accumulates at the TSS and further into the 5’ ends of gene bodies, however, it does also suggest that additional, quantitatively minor, trends of mutation risk heterogeneity within gene bodies may exist (see below).

The PC1 of the gene body mutation rates is characterized by a lowered mutation burden mainly from mutational signatures SBS1, SBS15 and SBS10b (Fig. 1A, C). Each of these signatures contains a significant N<u>C</u>G>T component in its trinucleotide spectrum, suggesting and association with DNA methylation (Fig. 1C and Extended Data Fig. 1B-D). Moreover, signature SBS1 is considered a stereotypical mutational process arising at methylated cytosines in a CpG context, presumably due to spontaneous deamination^36,37^. This localization strongly correlates with the average DNA methylation levels at those positions (Fig. 1E and Extended Data Fig. 1B). An association with DNA methylation also fits with the difference we observed between gene expression bins, where higher expressed genes (expected to have lower DNA methylation at promoters) showed higher weights of PC1 in SBS1 and other NCG-rich mutational signatures (Fig. 1C and Extended Data Fig. 1D).

The third mutation rate trend, PC3, explains less variance (∼3%), (Fig. 1D-F and Extended Data Fig. 1E) and its profile did not correlate with local DNA methylation profiles (Extended Data Fig. 1B), suggesting it originates from a different mechanism. This component is characterized by a sharp, hotspot-like mutation risk increase directly upstream of the TSS. Among the mutation signatures and tissues that contribute highly (Extended Data Fig. 1E and Supp. Fig. 2, 3) are UV damage in skin tumors (SBS7a and SBS38), CTCF-site enriched processes in gastric cancers (SBS17a) and, mutations in lymphoid blood cancers (Supp. Fig. 3). The localization of the peak as well as these SBS and tissue associations in PC3 are consistent with the known promoter hotspots at ETS transcription factor binding sites due to UV damage^19,20^, and CTCF-binding site hotspots associated with SBS17 mutagenesis^16^, as well as promoter-targeted SHM in lymphocytes^48^.

Overall, the trends in local mutation risk at the gene scale that we identified via PC1 and PC2 explain considerably more variation than the known promoter-associated hypermutation processes summarized in PC3. Further PCs PC4 and PC5 showed less clear patterns across the gene body (Extended Data Fig. 1F) suggesting that any further trends in mutation risk variation across gene bodies can have only modest effect sizes, compared to the PC1-PC3.

To aid interpretation, we further characterized the variability in the input matrix with a sparse PCA (see Online Methods). This yielded two clear patterns, (Extended Data Fig. 1G, H), recapitulating the mutation rate variation near the TSS, and independently a variation that extends within the gene body for at least 2 kilobases (Extended Data Fig. 1H, I). Among the contributing signatures, SBS9 and SBS37 present an enriched T>C spectrum, consistent with the error-prone polymerase POLH activity in the somatic hypermutation process.

Overall, our analysis suggests that DNA methylation is the main contributor to mutation rate variability along human gene bodies in multiple tissues and across gene expression levels. There is a substantial reduction of mutation rates at the TSS and extending downstream into 5’ ends of gene bodies, primarily affecting mutational signatures characterized by a CpG dinucleotide spectra.

### Gene stratification by local DNA methylation profile reveals epigenomic signature of intragenic enhancers

We hypothesized that different groups of genes, for instance due to varying expression level^36,49^, would show distinct shapes of local DNA methylation profiles and thus also varying shapes of mutation risk gradients along their gene body. To test this, we used the whole genome bisulfite sequencing (WGBS) DNA methylation data averaged along multiple solid tissues (see Methods and Supp. Table 2) to profile the DNA methylation levels along each gene body. In brief, gene bodies were segmented into 50bp bins, extending from the TSS and the TES inwards (i.e. towards gene center) by 5kb and extending outward (i.e. gene flanking regions) by 3kb, and the DNA methylation level averaged within every bin. The resulting profiles were then factorized using a PCA (see Methods, Fig. 2A and Extended Data Fig. 2A). Expectedly, the resulting DNA methylation profile PCs (mePCs) correlated with gene expression (Fig. 2B and Extended Data Fig. 2B). We considered the first three mePCs (together accounting for 27% of the systematic variability), representing the DNA methylation levels globally in the gene body (mePC1), the local TSS-proximal methylation level (mePC2) and the intergenic upstream and downstream methylation levels outside the gene (mePC3) (Fig. 2C). These were used to cluster human genes into 5 types, based on their local DNA methylation profiles (Fig. 2D, E and Supp. Table 3).

**Fig. 2.**
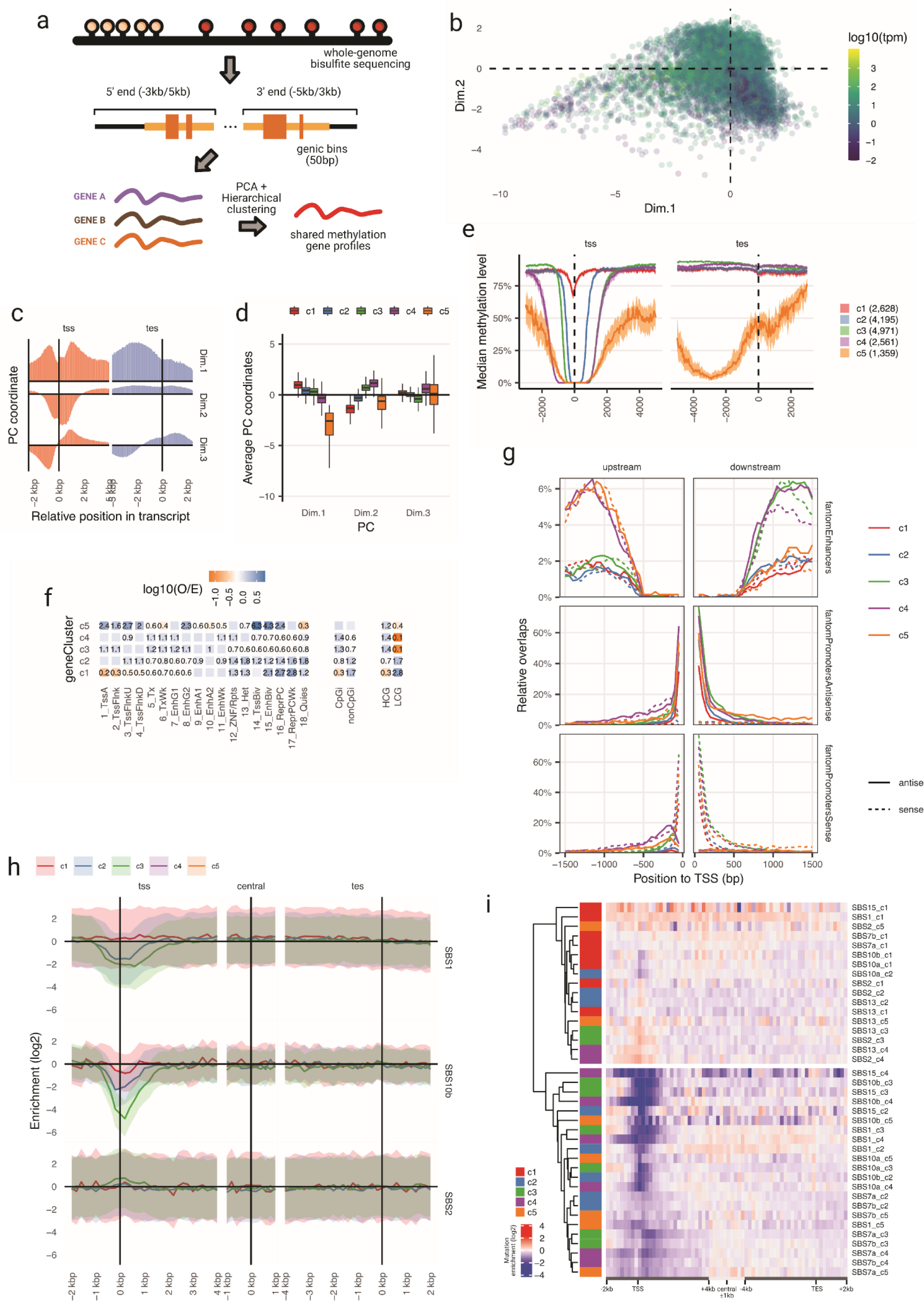
Clustering of genes according to their methylation profile. (a) Diagram of the gene clustering based on methylation gradients. Genes are tiled with 50bp bins and the average methylation value is calculated per bin. Gradients are then put together into a PCA and clustered using hierarchal clustering. (b) PCA coordinates of each gene from the factorization of methylation profiles. Color represents average gene expression across tissues. (c) PCA weights for the three first components used in the clustering of the methylation profiles. (d) PCA coordinate distribution of each gene cluster for the first three principal components. (e) Median methylation level for all genes in each cluster. Area represent the 95% confidence interval of the median across all genes in each group. (f) Overlap enrichment measured from observed and expected counts from a contingency table. Significant values are shown as numbers. Colors represent the logarithm in base 10 of the O/E score. Numeric values represent the untransformed O/E value. (g) Profiles of transcriptional data measured by CAGE from the FANTOM consortia53 and subdivided across promoters, enhancers and sense and antisense transcription (h) Mutation rate gradients of a selected set of mutational signatures (SBS1, SBS10b and SBS2) in a selected set of methylation profile gene groups. (i) Same as in (h) but with all gene groups for a wider set of signatures.

These 5 gene clusters exhibited distinct epigenetic characteristics (Fig. 2F and Extended Data Fig. 2C, D). Group 1 (c1), and to some extent group 2 (c2), contained genes with a methylated promoter (Fig. 2E), lower expression (Extended Data Fig. 2D) and a depletion in active transcription chromatin states (see Online Methods and Fig. 2F). In addition, the overlap with CpG islands^50^ and CpG-rich promoters^51^ was strongly depleted in these lowly active gene groups c1 and c2 (Extended Data Fig. 2C). The differentiating factor of c1 versus c2 groups was that c2 shows hypomethylation at promoters, resembling more active genes (Fig. 2F).

In contrast, gene methylation group 3 (c3) and 4 (c4) each represent a set of active, expressed genes (Extended Data Fig. 2D) with a wider hypomethylated section that encompasses the promoter. Importantly, we note also that this hypomethylation extends into the 5’ end of the gene body, approximately 2kb downstream of TSS, for both the groups c3 and c4 (Fig. 2E). While hypomethylation of DNA at gene promoters is well-known, it is less clear what generates the hypomethylated regions extending into gene bodies. Because these gene groups, c3 and c4, overlapped with active enhancer chromatin states (Fig. 2F), we hypothesized that activity intragenic enhancers could be one mechanism generating local hypomethylation within gene bodies. To test this, we analyzed nascent RNA transcription (measured by Cap Analysis of Gene Expression, CAGE)^52–54^, suggestive of enhancer activity. This has indeed shown CAGE signal in their corresponding 5’ gene body hypomethylated section of the c3 and c4 active genes (Fig. 2G). Of note, the intragenic enhancer transcription was detected in both DNA strands (Fig. 2G, dashed versus full lines), which is consistent with the activity of an active intragenic enhancer^54^. The main difference between c3 and c4 groups is the extent of the unmethylated section: c3 only presents the hypomethylation downstream of TSS into the gene body, while c4 has an extended hypomethylated region both upstream and downstream with similar size (Fig. 2E). Interestingly, the enhancer RNA transcription follows the same positioning, with c3 showing higher CAGE signal only in the TSS downstream section of the 5’ gene body, but c4 in both directions away from TSS (Fig. 2G). Further characterization of the methylation differences between the gene groups revealed that the CpG island shores^55^, regions adjacent to the island, could broadly account for these differences between c3 and c4 group (Extended Data Fig. 2E).

The gene group 5 (c5) contained generally shorter genes (Extended Data Fig. 2D) with, atypically, an overall less methylated gene body across its whole length (Fig. 2E and Supp. Fig. 5). These genes overlapped with repressed TSS and enhancers states (Fig. 2F) and were, consistently, enriched with the Polycomb gene silencing mark H3K27me3 (Extended Data Fig. 2G). Most of the homeobox genes, developmental genes with roles in cancer that were reported to be hypomethylated^56^, were preferentially included in this group (representing more than ∼7.5% of the c5 genes, Extended Data Fig. 2F). Polycomb repressive mark was previously associated with longer DNA hypomethylated segments. referred to “canyons” and “valleys”^57,58^, consistent with patterns observed in our gene group c5.

The averaged histone profiles of each gene category (Extended Data Fig. 2G) revealed an enrichment of the promoter mark H3K4me3, the enhancer/promoter mark H3K27ac and the enhancer mark H3K4me1, particularly, for clusters c3, c4 and c5, while this was modest or nonexistent for c1 and c2. Next, we asked whether the differential presence of these chromatin marks could support the mechanism where hypomethylation within gene bodies is causally linked to active intragenic enhancers. By performing a regression analysis, we found that the active transcription initiation features however also enhancer-like chromatin features (the H3K4me1 mark within the gene body) are particularly relevant for the distinction of the gene groups with different DNA methylation profiles (Extended Data Fig. 2H). Combing the histone modification profiles with the nascent enhancer RNA transcription described above, the existence of intragenic enhancers thus explains some cases of the unmethylated region extending downstream of the TSS and into the 5’ part of the gene body. We note that this enrichment of enhancer features towards the 5’ of gene bodies represents an average trend in many genes but in individual examples intragenic enhancers may be located at various positions in the gene. Previous associations of higher gene expression with lower DNA methylation levels specifically in the first exon and first intron, as well as patterns of evolutionary conservation, are consistent with a preferential 5’ gene body placement of intragenic enhancers^59,60^.

### Sub-gene scale mutation rate gradients differ across DNA methylation-based gene groups

Upon stratifying genes by local DNA methylation profile shape, we repeated the local mutation rate analysis to ascertain the role of DNA methylation changes in the local, sub-gene mutation rate gradients. In other words, we asked if the mutation risk gradients change between genes as the DNA methylation gradients change; if so this would further support the causal role of DNA methylation in mutation risk. We also considered mutational signatures separately to test for consistency across mechanisms. The genes having DNA hypomethylation at and/or nearby their promoters – those in groups c2, and the active gene groups c3 and c4 – showed a stronger depletion of mutational signatures depleted at 5’ gene ends, mainly SBS1, SBS10b, and SBS15 (Fig. 2H). Conversely, those mutation signatures that were enriched at the 5’ gene end, (APOBEC SBS2 and SBS13 and the AID-associated SBS9), showed an increased rate around promoters in gene clusters c3 and c4 (Fig. 1B, Fig. 2H, I). In contrast, in the c1 group mutation rate was constant across gene bodies, in accord with the rather homogeneous DNA methylation levels across their 5’ gene part (Fig. 2I). Genes in the cluster 5 (c5) group had similar profiles as the active genes c3 and c4, (Fig. 2I) but presented a less localized hypomutation, consistent with the hypomethylation of the whole gene locus.

Further, we considered a possible association between these gradients in DNA methylation, and the mutation risk in comparing exonic *versus* intronic DNA. Because of the enrichment of exons towards gene starts and ends (by definition of the gene model), and in light of reports of subtly different mutation rates ^61–63^ and subtly different DNA methylation in exons versus introns^64,65^, we considered our comprehensive WGBS dataset analyses. We checked the DNA methylation level of exons and introns, separately for each exon/intron in sequence, for a representative gene set (middle tertile of genes by length, and middle tertile in expression level). The resulting profiles showed that while methylation in the first exon was substantially lower compared to the first intron, consistent with the exon’s more 5’ positioning, the DNA methylation levels across the subsequent introns and exons were highly similar (Supp. Fig. 5C). Thus, in human WGBS data, after accounting for 5’ gene end hypomethylation, we see no notably different DNA methylation in the exonic versus intronic loci, and any intron/exon differences in mutation rates do not stem from different DNA methylation.

Overall, this mutational analysis of genes stratified by DNA methylation profiles provides evidence that the main determinant of the mutation rate variability along human gene bodies is local DNA methylation, which however has different repercussions on different genes and mutational processes. More highly expressed genes, and genes with intragenic enhancers, have more prominent and wider mutational coldspots toward their 5’ ends, respectively, when considering common mutational processes such as aging-associated SBS1, and some DNA repair failures (SBS15 and SBS10b). These trends are however reversed for APOBEC/AID mutagenic signatures, which are enriched at hypomethylated DNA found at active promoters and intragenic enhancers.

### Lowly methylated regions within or without genes show consistent hypomutation

Following up on the analysis of gene body mutation rates, we hypothesized that DNA methylation has roles in shaping local mutation rates in various loci across the genome, even if not contained within genes. To test this, we identified a dataset of genomic segments that are uniformly hypomethylated in various cell types, either with a near-complete lack of DNA methylation, unmethylated regions (UMRs) on in lowly-methylated regions (LMR), with intermediate methylation levels^66,67^ (Fig. 3A).

**Fig. 3.**
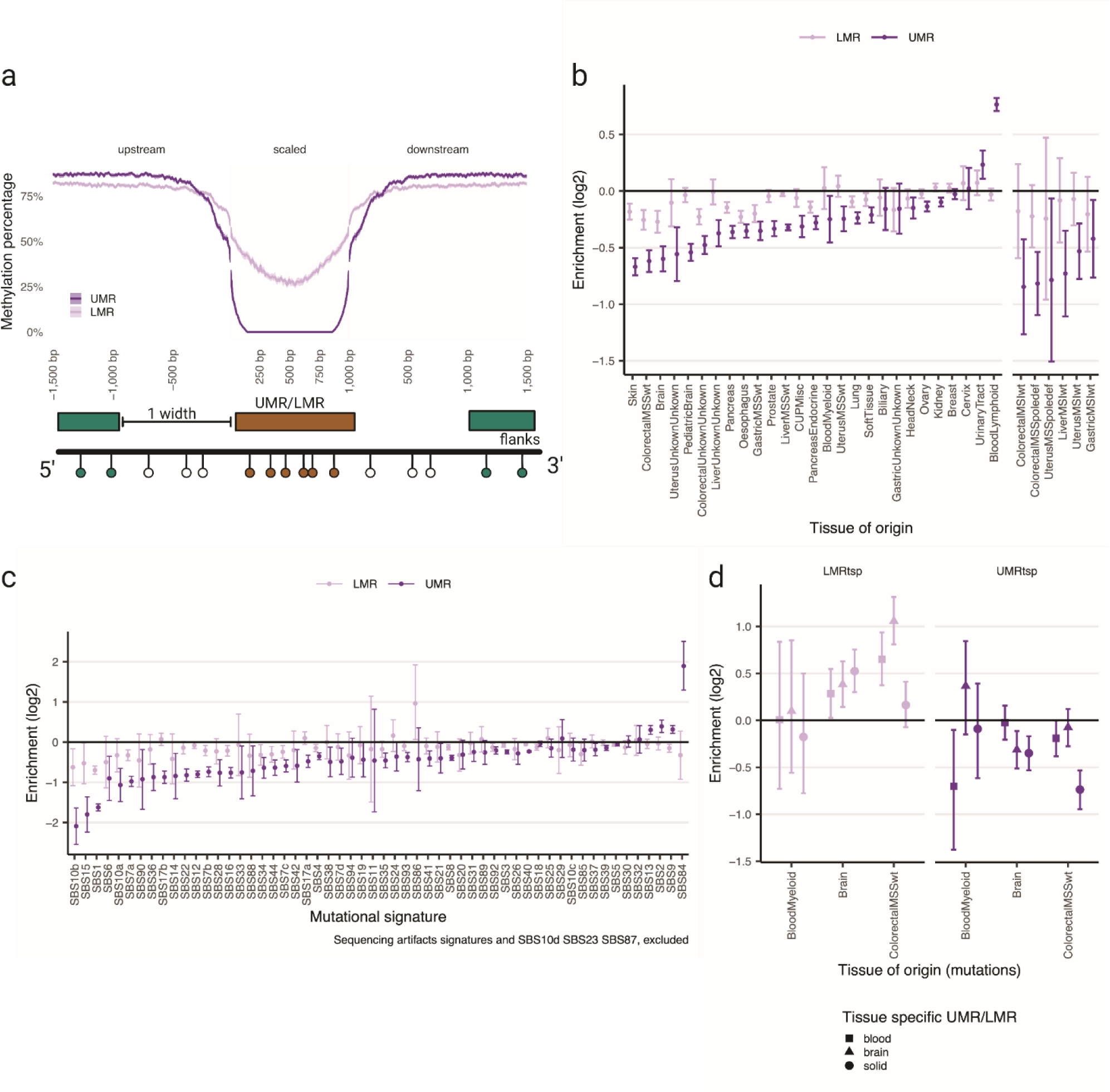
Mutation rate gradients in genome-wide hypomethylated regions. (a) Methylation profiles, measured as the median methylation level in each bin, for both UMRs and LMRs. Shadow area represents the 95% confidence interval of the median value across all regions. (b) Coefficients representing the relative mutation rate change for the UMR or LMR regions versus flanks. Each regression includes all mutations for a given tissue. (c) Same as in e but for the assigned mutational signatures. (d) Coefficients measuring the relative mutation rate change in tissue specific UMRs and LMRs versus flanks.

Firstly, we curated a set of hypomethylated regions in the human genome from previous publications^56,67^ (Supp. Table 4), which contained 18 tissues and represented predominantly stem cells and blood cell lines. Additionally, we collected genome-wide (WGBS) DNA methylation data from public repositories (Roadmap^68^ and Encode^69^) and called UMR and LMR loci using MethylSeeker^67^ (see Online Methods and Supp. Table 2). Here, we considered 34 diverse solid tissues (non-neural) plus 6 blood, and 4 brain tissues, a selection that represents better the sequenced tumors we analyzed (see Online Methods; the solid, blood and brain tissue groups are treated separately because of considerable differences of epigenomes between them^69^). In total, all obtained UMRs spanned 41Mbp, similar to previous reports, while LMRs detected in this study covered a total of 25 Mb (Extended Data Fig. 3A, B).

Most of the surveyed tissues, except the urinary tract cancers (often very highly mutagenized by APOBEC) and the lymphatic blood cancers (often showing evidence of somatic hypermutation via AID), showed a significant overall reduction of the mutation rate at UMRs and LMRs, with an average depletion across tissues of ∼25% (Fig. 3B). This local hypomutation was more substantial for tissues with a high proportion of SBS1 mutations, like colon and brain^8^. Skin cancers also showed a significant reduction in mutation rate at UMRs, consistent with a proposed role of DNA methylation in the predisposition to UV damage mutations (Fig. 3B)^70^. These associations were highly consistent when tested on either the set of UMRs obtained from previous publications, or the ones computed in this study (Extended Data Fig. 3C).

Considering individual mutational signatures, where mutations were pooled across tissues in a pan-cancer setting, mutational signatures SBS1, 10b and 15 decreased the most in UMRs, which is consistent with our analysis of hypomethylated 5’ gene ends above. UMRs had on average ∼75%, ∼65% and ∼55% fewer mutations than expected by trinucleotide composition, for SBS10b, 1 and 15, respectively. Other mutational signatures like SBS6, related to MMR deficiency, and SBS7a resulting from UV exposure also showed a notable reduction of mutations at UMRs (Fig. 3C).

Again consistently with gene body analysis, certain signatures showed an increased mutation rate at UMRs genome-wide. SBS84, associated with the activity of the activation-induced cytidine deaminase (AID) in the SHM process^71^ in lymphocytes showed a four-fold (2.5-5.6 fold; 95% CI) increase, consistent with AID acting at or nearby highly active gene promoters^48^. Consistently, SBS9, also associated with SHM in lymphoid tissues in part reflecting the activity of DNA polymerase η, did so as well to a lesser extent (29%). Finally, the widespread signatures SBS2 and SBS13 resulting from APOBEC mutagenesis also showed an enrichment in the UMRs (35% increase over expected mutation rates for SBS2 and 28% for SBS13) (Fig. 3C). Thus, AID and APOBECs preferentially mutate hypomethlyated DNA, which is common at active gene regulatory elements.

### Tissue-specificity in hypomethylation-associated mutational coldspots

We considered the differences in local DNA methylation between tissues, testing their association with local mutagenesis, to provide evidence for causal roles of DNA methylation in mutation rate changes. Here, we examined a set of UMRs and LMRs from each tissue group and compared it to the relative mutation rates in matched *versus* mismatched cancer types. As representatives from the solid tissue group, we chose three datasets from the digestive tract, comparing them with the colorectal cancer mutations. Of note, while UMRs showed a high consistency across tissues, LMRs were more tissue specific (Extended Data Fig. 3D, E), consistent with the LMR-associated genomic features, i.e. enhancers^66^. The total yield of tissue-specific UMRs was thus sparse (Extended Data Fig. 3E). In the selected sites, the depletion of mutations in UMRs from the matching tissue was more evident than in the UMRs of the mismatched tissues. For instance, colon cancers showed a 40% lower mutation rate for UMRs specific to digestive tissues, while the reduction of mutations in brain and blood cancers at digestive tissue UMRs was not as substantial. Similarly, the reduction of mutation rates in the blood-specific UMRs was 37% (7% – 61%, 95% CI) in myeloid blood tumors but only 4% (−30% – +35%, 95% CI) depletion for colon cancers. Our brain enriched UMRs showed a less striking selectivity (Fig. 3D).

Overall, the observed variability in local mutation rates at UMRs at the tissue level is explained both by to what signatures the tissue is normally exposed and additionally by the tissue-specific variation in occurrence of hypomethylated loci.

### Variable mutation rates at UMRs that intersect important functional genomic elements

Due to the known characteristic hypomethylation of multiple regulatory elements like promoters, enhancers and CTCF/cohesin binding sites^72–75^ (here, we considered chromatin loop anchor points from cohesin ChIA-PET experiments^76^) we used these annotations to classify the extracted UMR sets. This was to ask whether the methylation effect on mutation rate is particular to some functional elements or is a general property of DNA hypomethylation also seen outside known functional elements (Fig. 4A).

**Fig. 4.**
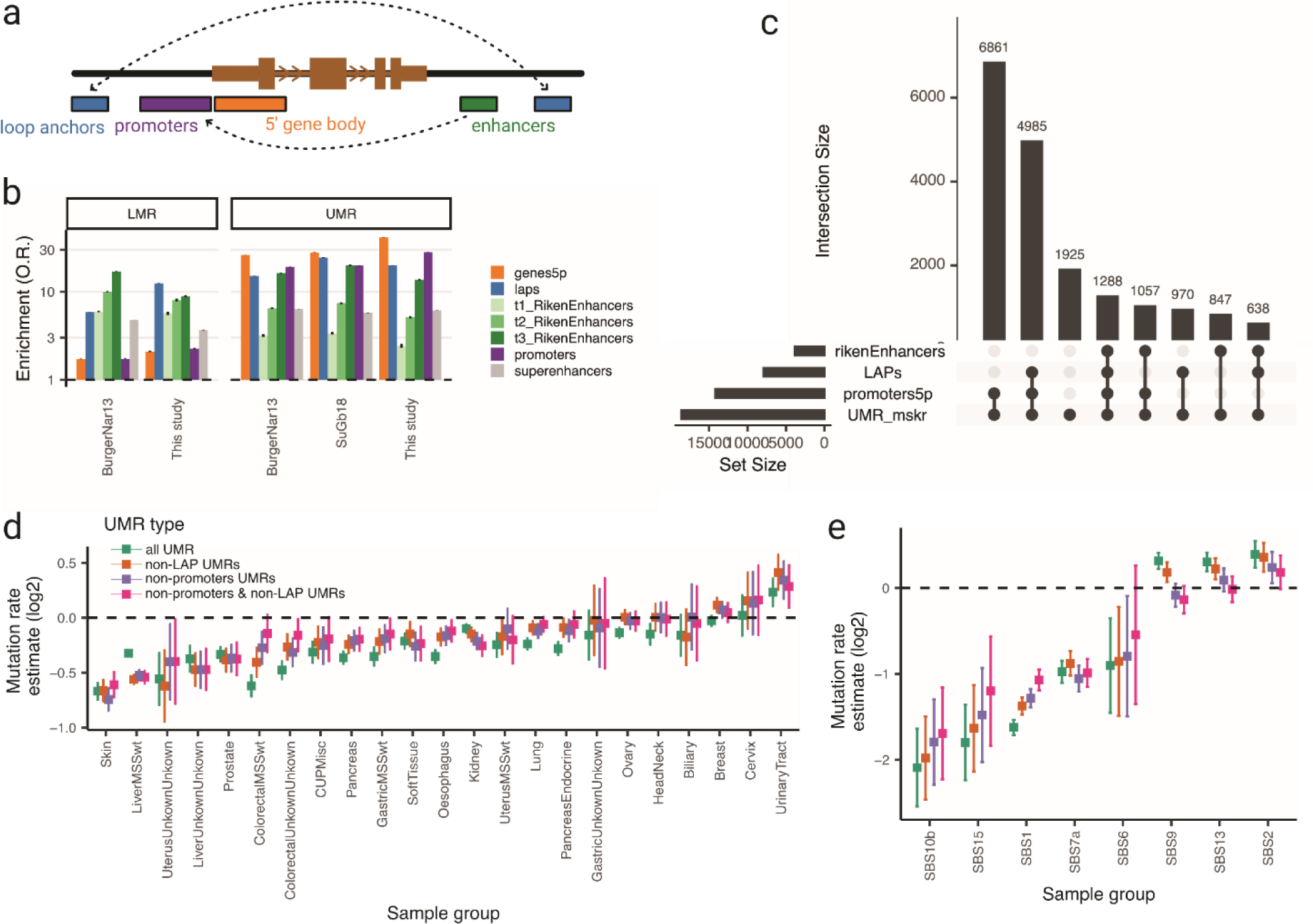
UMRs and interactions with other functional elements. (a) Diagram of the set of the relevant functional elements represented here. (b) Odds ratio enrichment of the overlap of a given functional element either with the UMR or the LMR. (c) Upset plot showing all possible intersections of UMRs with the functional elements depicted in (a). In this panel, the 5’ end of the gene body and the promoter is mixed in a single group. (d) Mutation rate enrichment for UMRs that do not present an overlap with functional promoters or loop anchors. (e) Same as in (d) but for the stratified mutational signatures.

As expected, UMRs were enriched in promoters as defined at the TSS upstream region, and LMRs in enhancers, as defined by CAGE experiments from the FANTOM dataset^53^ (Fig. 4B). This held true for both prior UMR sets and the UMRs called in this study, and so being consistent between tissues and methodologies. The highest number of UMRs was explained by promoters and the promoter-adjacent 5’ ends of gene bodies (37% uniquely and 76% explained together with other elements) (Fig. 4C). However, 10% of the UMRs genome-wide did not overlap with the above-mentioned known functional elements (Fig. 4C) and for LMRs, this was up to 52% (Extended Data Fig. 4A), broadly consistent with previous estimates of UMR and LMR overlap with functional elements^66^.

We then asked if the local reduction in mutation rate was, at least in part, due to some other property of these functional elements (promoters, enhancers or CTCF/cohesin bound loop anchors), rather than necessarily their DNA hypomethylation itself. For every tissue, we removed the UMRs that overlapped either the promoter adjacent region, defined as 2kb upstream and 1kb upstream of TSS, and chromatin loop anchors from the UMR set of interest. (Note that enhancers were not considered in this analysis because of little overlap with UMRs). The overall trend of hypomutation was still evident in remaining UMRs, both across tissues (Fig. 4D and Extended Data Fig 4B) and signatures (Fig. 4E), albeit at somewhat lower magnitude. As an example, SBS1 was hypomutated 69% in the full UMR set, and hypomutated to a similar level (51%) in the UMRs not overlapping promoters or cohesin loop anchors; SBS15 depletion was reduced slightly from 70% (all UMRs) to 54% (UMRs outside known elements). We note that the UMR sets that overlap with promoters and loop anchor points did show a slightly stronger hypomethylation (Extended Data Fig. 4C).

Thus the mutational effect at DNA hypomethylation loci is largely independent of their overlap with known functional elements. This implies that some additional mutation coldspots resulting from hypomethylation can occur elsewhere in the genome, unrelated with known promoters or enhancers or loop anchors. We do not exclude that this results from insufficient power to thoroughly detect additional enhancers or loop anchors, which may in some cases have a high degree of cell type specificity, using the data in our analysis.

### Hypomutation of functional elements is largely explained by hypomethylation thereof

The above analysis suggests that hypomethylation is causal to local mutation rate reduction at functional elements genome-wide. To further examine if hypomethylation is the main determinant of local mutation risk in regulatory DNA, we examined if the promoters and the CTCF/cohesin-bound chromatin loop anchors exhibit any additional mutation rate reduction, after accounting for their hypomethylation.

As anticipated from their overlap with identified UMRs, the median methylation value around promoters and CTCF/cohesin chromatin loop anchors shows a substantial depletion (Extended Data Fig. 4D, E) of the average methylation values.

While the hypomethylation of CTCF bound sites is known^49,73^, we note that in the broader region containing the chromatin loop anchors (here from cohesin ChIA-PET^76^), the methylation levels were reduced suggesting hypomethylation extends further than the CTCF binding site. This hypomethylation at loop anchors is consistent with the overlap between promoters, UMRs and CTCF binding sites (Extended Data Fig. 4E), but appears not fully explained thereby, suggesting that chromatin loop anchor regions are hypomethylated, regardless of their spanning promoters or included CTCF sites.

The mutation rate (relative to flanking DNA) was reduced at both promoters and chromatin loop anchors, and importantly this reduction depended on DNA hypomethylation (Extended Data Fig. 4F). The promoters that did not overlap sufficiently with an UMR did not show appreciable hypomutation (Extended Data Fig. 4F). SBS1 mutations showed an overall 51% depletion in all chromatin loop anchors, while showing a lesser 30% depletion in those loop anchors that did not contain a detected UMR, and a similar depletion was also seen in other methylation-associated signatures like SBS15 and SBS10b. Of note, the reduction of mutations in neither promoters nor chromatin loop anchors was as striking as when measuring the UMR itself, possibly due to only a partial coverage of annotated functional site with the unmethylated loci. This is consistent with a leading role of DNA hypomethylation in determining local mutation rates in regulatory DNA genome-wide. In addition, other known factors at these regions such as chromatin accessibility^23^ may further modulate mutation rates^24^.

To rule out that the effect of the UMR on mutation rates was indirect and resulted from the increased transcription of genes with a UMR, we repeated this analysis after stratifying genes by mRNA expression (Extended Data Fig. 4G). For colorectal and skin tissues, which contained sufficient mutation counts, the local mutation rate in genes with high mRNA expression but without overlapping UMR was not reduced, suggesting transcription levels do not explain the mutation rate decrease. The relative mutation rate of gene-overlapping or adjacent UMRs in the middle and high - expressed bins (Eq2 and Eq3) was equivalent and significantly reduced compared to their UMR-less counterparts of same expression level (Extended Data Fig. 4G). Of note, some tissues like liver (Extended Data Fig. 4G) did show reduction of mutation rate in higher expression bins, consistent with known roles of transcription-coupled processes, however even so, mutation rates were still reduced in UMR-overlapping promoters (Extended Data Fig. 4G).

In summary, DNA hypomethylation determines local mutation rates in a manner independent of other features at regulatory elements and chromatin loop anchors and independent of transcription levels.

### Local methylation-aware mutation rate baselines can clarify signatures of selection

Methods to detect selection on somatic mutations in protein-coding genes and other functional elements rely on an accurate baseline of local mutation rates. This is used to test whether there is an excess or dearth of mutations over that baseline, signifying positive or negative selection, respectively. Such baselines are often established at the whole-gene level, and are based on a variety of mutation rate covariates such as DNA replication time, mRNA expression levels and histone marks.

DNA methylation profiles of genes or other functional elements, capturing local variation in DNA methylation, may be considered in order to establish more precise baselines for mutation rates that account for the within-gene heterogeneity in mutation rates. To provide a proof-of-concept analysis for utility of DNA methylation-aware baselines for mutation rates, modeled the mutation burden of each gene from the 10,295 TCGA whole exome sequences^77^. Because mutation rates are heavily influenced by the epigenetic state and the replication time^78^, we predicted gene mutation rates from a set of existing epigenomic covariates from a state-of-the-art tool (dNdScv)^12^ as a base model. We then compared this with a model that additionally includes the DNA methylation-profile gene groups labels defined above (c1-c5) as an additional covariate. As a negative control, we consider the same model but with the DNA methylation group labels shuffled (Fig. 5A and Methods). The root mean square error (RMSE) in number of exonic (including UTRs) mutations per gene, using the average of 15 runs of five-fold cross validation, showed a better fit for the DNA methylation-aware model compared to both the base (covariate-only) model, and the shuffled methylation-covariate model (Fig. 5B). Significance of this improvement was tested by comparing the deviance using the observed *versus* DNA methylation-shuffled group labels (see Online Methods). While in all iterations in the DNA methylation-aware model, the feature had a significant effect on improving the fit (p-value always < 2e-16 in all 75 iterations), the shuffled-methylation label gene groups models were significant in only <10% of the crossvalidation iterations (6 out of 75 at p-value < 0.01) (Extended Data Fig. 5A).

**Fig. 5.**
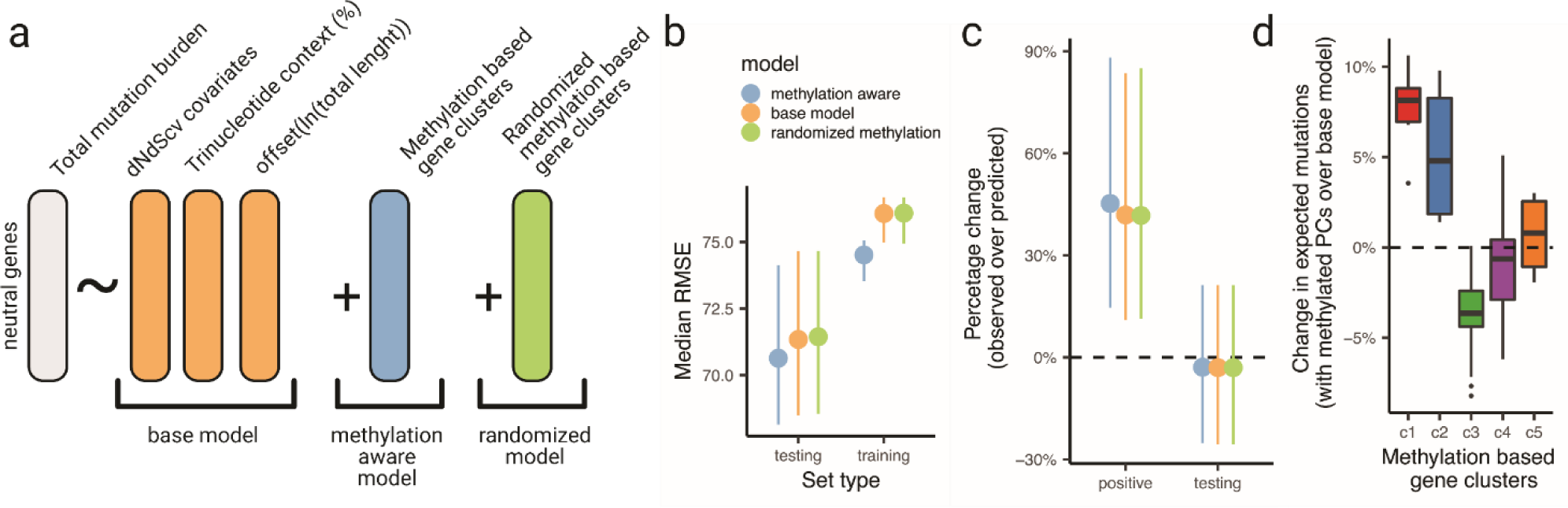
Methylation aware groups are relevant for selection estimation. (a) Diagram depicting a model to predict the mutation rate of genes according to epigenetic covariates12, the context composition of the gene and the length as a offset. To this base model, the methylation-aware gene classes are added together with a randomized version of the gene clusters. (b) Root mean square error for the prediction of mutation rates by each model. (c) Percentage change of predicted mutations in the positive set (cancer genes with positive selection) and the testing set (genes that are used to evaluate the performance of each cross-validation round). (d) Changes in the predicted mutations burdens of genes for each gene cluster as defined in Fig. 4.

Using the expected number of mutations predicted by this model, we can calculate the excess mutation burden for every gene, which is an effect size estimate of positive selection. As expected, this mutation burden excess was on average 45% in known broadly-acting known cancer driver genes (i.e. drivers in more than 2 tissues according to MutPanning^79^, n= 131), which were excluded from the training and testing partition of the original data (Fig. 5C). When considering non-cancer genes there was no significant excess in mutation rates between the different models, DNA methylation included as a covariate or not (Fig. 5C).

Further indicating that gene DNA methylation patterns are an important factor to account for in identifying positive selection, the expected mutation burden differed significantly between different methylation-profile gene groups (Fig. 5D and Extended Data Fig. 5B). In particular, genes in the c3 group corrected their mutational burden prediction to lower values upon including DNA methylation covariate (with a −4% median adjustment in considered set of tumor, although this may differ in other datasets depending on the active trinucleotide mutational signatures). Example genes in this group are *EGFR*, *KRAS* or *RB1*, where the corrected expectation (to lower values) raised the observed excess mutation burden, suggesting a stronger positive selection (Extended Data Fig. 5B). Conversely, genes in group c1 and c2 (with a narrower and/or weaker hypomethylation region) showed a correction towards higher expected burden values when considering DNA methylation (with a median of +5% mutation burden for c1 and +3% for c2). *TSC1, FAM135B* and *APC* are examples of these groups, which show an overall reduction in their excess burden, suggesting the positive selection estimates would be revised downwards upon considering DNA methylation (Extended Data Fig. 5B). The other methylation aware gene groups, c4 and c5, showed only modest corrections (<1% change in excess mutation burden; median within group, although individual genes may have bigger deviations). Of note, genes classified by the Cancer Gene Census as either oncogenes (44.1%) or tumor suppressors (45.1%) were enriched in the active hypomethylated group c3 (Extended Data Fig. 5C). Thus, our model can capture relevant information for determining the mutation burden estimates for genes stratified by their DNA methylation profile.

## Discussion

Somatic mutation rate variability was studied thoroughly at single nucleotide resolution, tallied across oligonucleotide sequences^8–10^, and it was additionally studied across megabase-sized chromosomal domains^1,7^. Prior work on the identification of mutation rate variability on the “mesoscale” in between these two extremes – the coarse-scale and fine-scale heterogeneity – has considered some genomic features that interact with damage formation or repair^14,23,80^. Here, motivated by the known intra-gene gradients in histone marks that tend to occur in active genes and that are relevant to DNA repair^30,31,35^, we systematically tested for the local mutation rate gradients along gene bodies across 40 mutational signatures, separately considering somatic tissues and gene expression levels. This highlighted the local DNA hypomethylation at the 5’ end of genes – at promoters and further extending ∼2 kb downstream into gene bodies – and more generally at other hypomethylated loci along the genome such as chromatin loop anchors, as a major influence on mutation rate heterogeneity in the human genome at the kilobase-scale.

In our unbiased analysis of mutation rates, which adjusted for trinucleotide composition of various gene segments, the 5’ gene part was the focus of the top trends in variation between mutational signatures and gene expression levels (Fig. 1D). The strong depletion of the common SBS1 “clock-like” signature was observed; in addition we noted the DNA repair deficiency-associated SBS10b and SBS15 (Fig. 1A, C), recently linked with DNA methylation^39,40^. Thus, the gradients in DNA methylation across genes cause locally variable mutagenesis, by generating mutational coldspots at 5’ ends of genes. An important exception where these coldspots change to hotspots are the APOBEC mutagenesis (SBS2 and SBS13) commonly observed across many somatic tissues, or the more specialized SHM associated signatures (SBS9, SBS84) observed largely in B-lymphocytes.

Similar to how mutations from the AID-initiated SHM process are well-known to be enriched at or near promoters^48^, we also show this is the case for 5’ gene end enrichment of the mutations by its APOBEC paralog(s), although the underlying mechanism may be different. The AID may sense 5’ stalled RNA polymerase^81^, while we speculate that APOBEC deamination may be more mutagenic at 5’ gene ends due to DNA methylation itself protecting against APOBEC in the remainder of gene body. Links between higher APOBEC activity and DNA demethylation have been reported^82^, however other studies did not show notable associations^83^. We note it is also possible that the causality flows the other way: rather than DNA methlyation blocking APOBEC, APOBEC could in principle remove DNA methylation in a non-mutagenic manner, instead of introducing mutations.

AID, a homolog of APOBEC enzymes, was claimed to participate in active DNA demethylation via triggering DNA repair^84,85^ and so, speculatively, an increased activity of APOBEC at 5’ ends, directed thereto by an unknown mechanism, might act to contribute to reduce DNA methylation; additionally the same mechanism would cause mutations in those regions.

We also show that a gene classification by the extent of their hypomethylated region at the 5’ end (Fig. 2E) is reflected in the extent and the intensity of the mutation coldspot in the gene 5’ section (Fig. 2H). This variable DNA hypomethylation and thus mutation rates within genes associates with occurrence of intragenic enhancer regions, chromatin loop anchors, or in some cases Polycomb marks, and possibly additional factors associated with hypomethylation yet to be identified. We suggest that gene body local DNA methylation variability should be included in models for testing selection on somatic mutations at the gene level. Finding the best implementation of this principle remains a direction for future work, where for instance selection on different gene segments might be considered individually. Further, methods that incorporate DNA methylation into mutation rate baseline may be developed for more accurately estimating selection on non-coding regulatory regions.

Prompted by the mutational coldspots in 5’ gene ends, we extended our analysis to genome-wide hypomethylated regions (both UMRs, and the partially methylated LMRs^66,67^). These were, as expected, associated with promoters and enhancers, however interestingly also with chromatin loop anchors suggesting hypomethylation is common at these loci (Fig. 4B, C). There was also a residual fraction of hypomethylated regions not overlapping known elements however still constituting mutational coldspots distributed across the genome, supporting anticipated mechanisms underlying mutation reduction at these sites^39,72^.

Overall, we highlight the role of local DNA methylation in shaping mutation rate heterogeneity in the human genome at the mesoscale and stress the need to further characterize the factors governing the local, sub-kilobase scale variation in mutation risk and the underlying mechanisms.

## Methods

### Mutation datasets used in this study

Somatic mutations used in this study were compiled from tumor-normal matched whole genome sequencing experiments available by prior studies (Supp. Table 1). In brief, single nucleotide variants (SNV) from a unified mutation calling dataset^86^ comprising the entire Pan-cancer Analysis of Whole Genomes (PCAWG^42^) were downloaded from the PCAWG resource site (https://dcc.icgc.org/releases/PCAWG/Hartwig) with ICGC application code DACO-1101. Somatic SNVs for a set of tumor and metastatic samples from the Personal Oncogenomics (POG) project^43^ were downloaded from BC Cancer, publicly available (www.bcgsc.ca/downloads/POG570).

Additional sets of SNVs from healthy tissue samples^44,45^ were compiled from the literature.

### Whole genome bisulfite data used in this study

Whole genome bisulfite sequencing data (WGBS) were downloaded from publicly available datasets, in brief, the ROADMAP^68^ epigenome project (see https://egg2.wustl.edu/roadmap/web_portal/) and the ENCODE^69^ data portal (see https://www.encodeproject.org/). From both sites, all available WGBS datasets were first selected and processed but only the ones that were found with sufficient quality and that passed manual inspection were used in the analysis (see Online Methods and Supp. Table 2).

Downloaded data consisted in fractional methylation and coverage files that contains information about the percentage of methylated reads of each sufficiently covered CpG and the original depth of that loci (Supp. Table 2). Additional sets of precomputed unmethylated and partially methylated genomic loci (UMRs and LMRs, see Online Methods) were also compiled from prior literature^56,66,67^ (Supp. Table 4).

### Factorization of mutation gradient profiles along genes

Genes were segmented in 250 bp bins along the gene 5’ and 3’ ends together with a central window that extended from the central position of the gene. The gene ends were expanded 2kb outward (upstream from 5’ end and downstream from 3’ end) and 4Kb inward. The central position was expanded 1kb in each direction. An extra 1kb reference bin was selected 4kb downstream from the 3’ end of the gene. Individual genes were grouped in 3 quantiles according to their average expression in the GTEX V8 portal^87^. The somatic mutations used in the analysis were first grouped according to the tissue and their assigned signature (Online Methods) and then intersected with the bins across the gene body, yielding a total of 512 sets. The mutational enrichments of each set were measured with a negative binomial regression and the resulting genic bin coefficients factorized in a single PCA.

### Gene classification from DNA methylation profiles

Genes were segmented similarly as for the mutational gradient example but at higher resolution of 50bp and without a central section. Both the TSS and the TES ends of genes were extended inward for 5kb and outward 3kb. Methylation averages (within 50bp segments) were computed using deeptools^88^. The resulting matrix contained an average methylation value per gene per segment. This matrix was factorized using a Principal Component Analysis (PCA) with no scaling (Online Methods). The resulting PC coordinates where clustered using medoids clustering, selecting 5 groups from a range of 2 to 7 after visual inspection of the genomic characterization and methylation gradients (Online Methods). To generate the methylation gradients depicted in this study (Fig. 2E), each segment in each gene was grouped according to its assigned cluster and the average value per segment was plotted and the 95% (two-tailed) confidence intervals were extracted from a binomial test at a given sample size equal to the amount of genes tested.

### Selection analysis

Selection in genes was estimated from whole exome data available from the mc3-TCGA dataset (Online Methods). For each gene we mapped the gene id to the epigenomic-based PC coordinates of dNdScv^12^ package to use as a base model in our predictions. We also extracted the trinucleotide composition of each gene and normalized them per gene. A negative binomial regression was used to predict the total number of mutations observed at a specific gene (the mutation burden). This approach was repeated three times, (1) with a base model containing the size of the gene, the trinucleotide composition and the dNdS cv covariates, (2) with the base model and the methylation aware gene grouping and (3) with the base model and a shuffled set of methylation groups that maintained the existing class imbalances of the original classification (Online Methods). Cancer associated genes were downloaded from mutpanning^79^ dataset with the added requirement that at least they must be associated with 2 cancer types and they were excluded from the initial fitting of the model.

## Acknowledgments

We thank Ahmed Khalil for assistance with retrieving datasets with gene models, and with somatic mutation data. D.M-P. was funded by a Severo Ochoa FPI fellowship (MCIU/Fondo Social Europeo; BES-2017-079820). F.S. was funded by the ICREA Research Professor program. Work in the lab of F.S. is supported by an ERC StG “HYPER-INSIGHT” (757700), Horizon2020 project “DECIDER” (965193), Spanish government project “REPAIRSCAPE”, CaixaResearch project “POTENT-IMMUNO” (HR22-00402), the SGR funding of the Catalan government, and the Severo Ochoa centers of excellence award of the Spanish government to the hosting institution. The authors acknowledge support from the Severo Ochoa Centre of Excellence program to IRB Barcelona. The results published here are in whole or part based on data generated by the TCGA Research Network (https://www.cancer.gov/tcga).

