## Extended Data Figures 1-5 for "Mutation rate heterogeneity at the sub-gene scale due to local DNA hypomethylation"

**Supplementary Material: Extended Data Figures 1-5** for Mas-Ponte and Supek (2023).

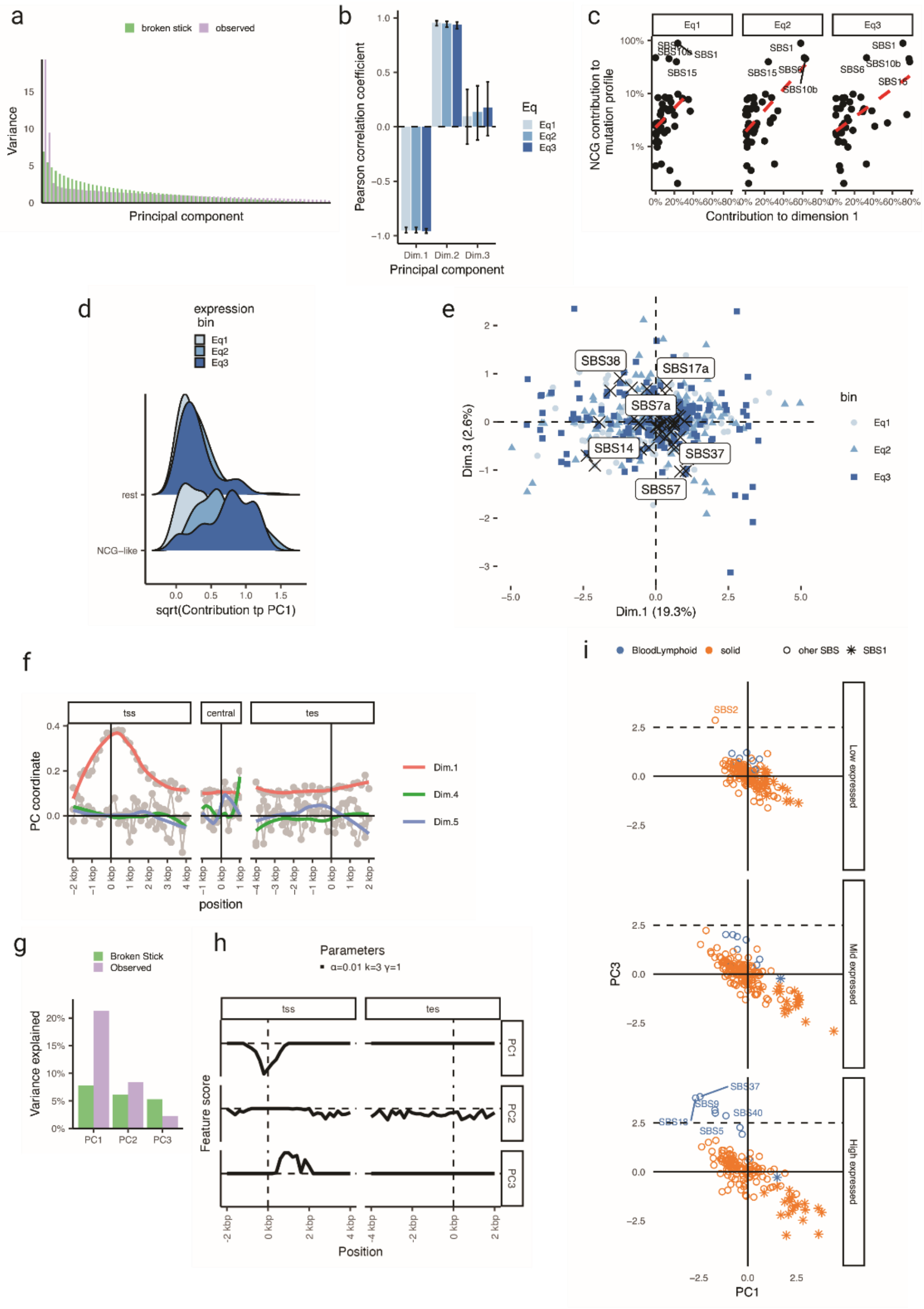

**Extended Data Fig. 1. Systematic quantification of mutation gradients along gene bodies.** (a) Scree plot from the gene gradient mutation rate PCA depicted in Fig.1. (b) Correlation of the percentage of CG trinucleotides in each signature compared to the total contribution to the first principal component. (c) Same as in (b) but instances are stratified by gene expression and signatures are classified in CG-like or rest according to the CG percentage in their profiles. (d) Signature contribution to PC1 as in (c) but grouping signatures according to high NCG (NCG-like) or low (rest) values. (e) Coordinates of mutation signature profiles in different sets, same as in Fig. 1C but with Dim.3. (f) Weight profiles of PC4 and PC5 (with PC1 as scale). (g) Scree plot for the sparse PCA performed in the mutational gradient matrix with 3 components. (h) Profiles of the 3 components extracted with a sparse PCA. (j) Coordinates of different signatures colored by tissue, blue for Blood Lymphoid samples and orange for other. Shape as an asterisk for SBS1 and a empty circle for other signatures.

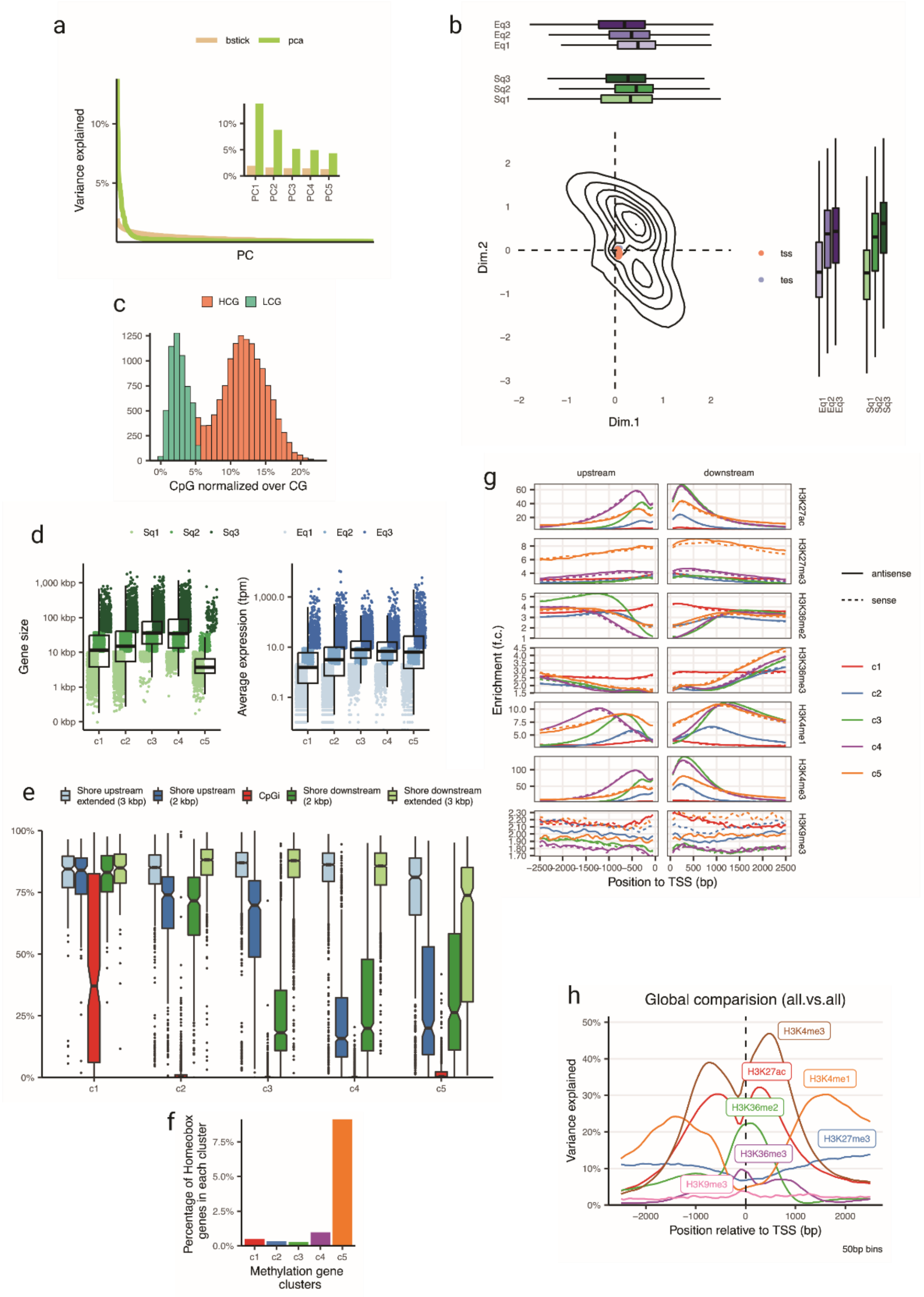

**Extended Data Fig. 2. Clustering of genes according to their methylation profile.** (a) Scree plot of the methylation profile PCA used to cluster genes. (b) PCA coordinates of each gene (represented as a 2D density plot) with the expression and size distribution for each principal component represented in boxplots. (c) Definition of the HCG genes according to their normalized CG values. A mixture modeling is used to define a numeric cut-off. (d) Expression and size terciles of each gene methylation class. (e) Enrichment of histone mark signal plots (fold change extracted from ENCODE ChIP-seq experiments) both for sense and antisense genes in each methylation aware group. (f) Proportion of Homeobox genes, as defined in ref[89], for each methylation aware group. (g) Methylation levels across different methylation aware gene groups including their CpG island shores (2kb from the upstream and downstream end of the CpG island) and a extended region (h) Histone weights per position (relative to e) in a linear model to classify methylation aware gene groups.

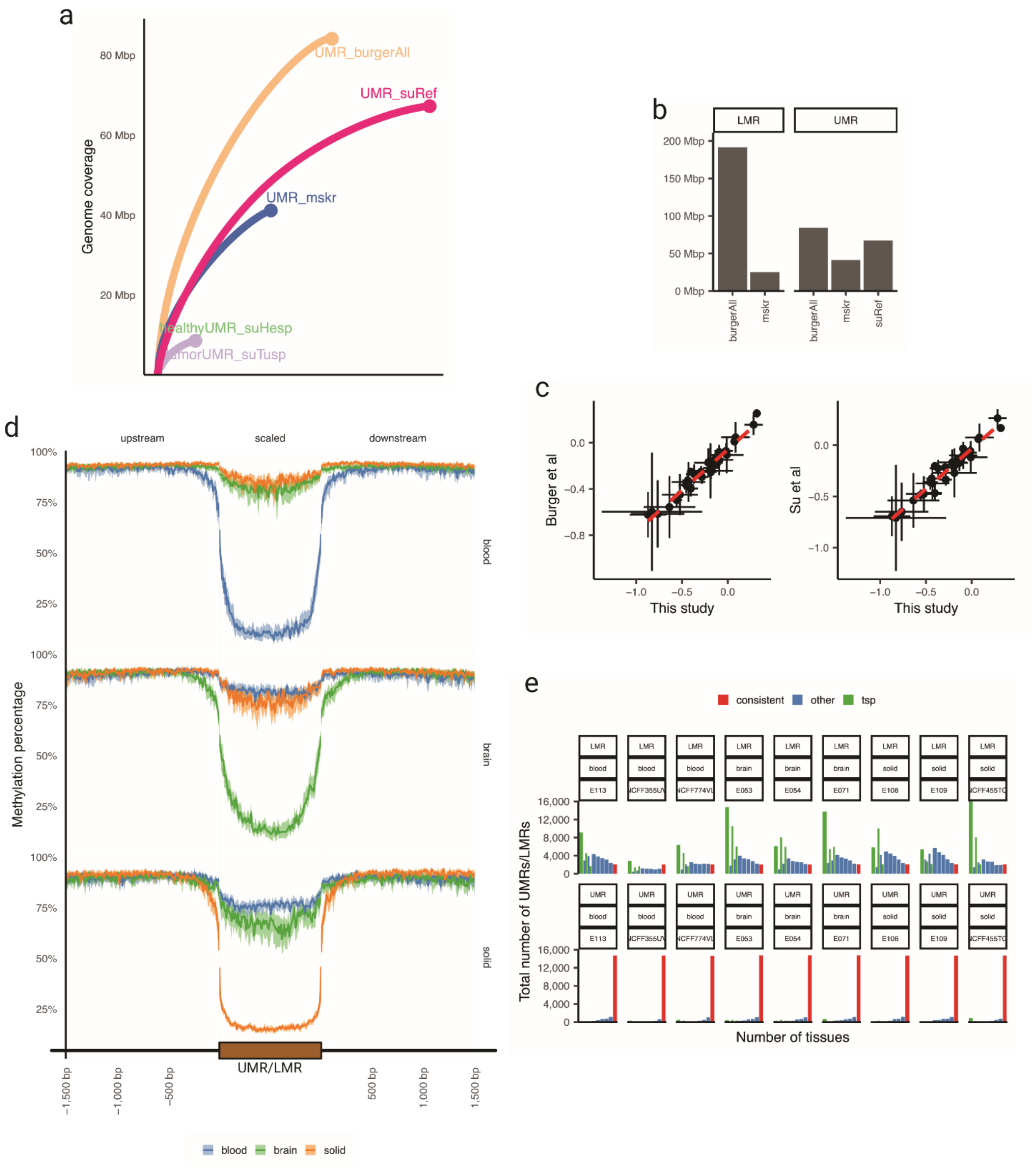


**Extended Data Fig. 3 UMR and LMR. Genome-wide unmethylated regions and quantification of mutation rate:** (a-b) Genomic coverage of the selected UMRs. (c) Correlation of the mutation rate estimations in different UMR sets. (d) Methylation levels in tissue specific UMRs. (e) Number of tissue specific (green), consistent (red) and other (blue) UMRs and LMRs from the comparison of 3 brain, blood and digestive datasets. In the x-axis, the number of sets that contain a given type of UMRs is represented, ranging from 1 to 9.

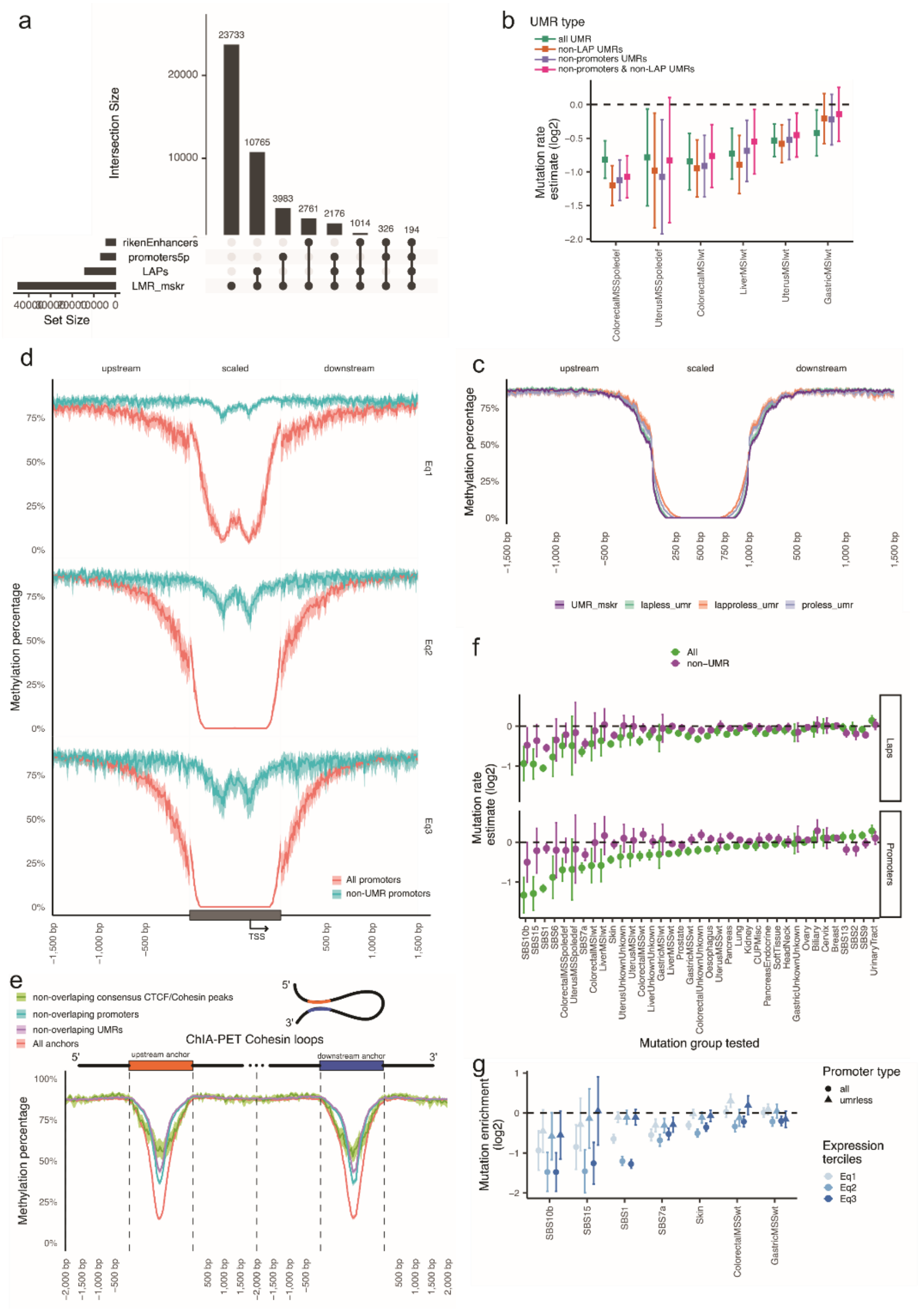

**Extended Data Fig. 4. Functional elements associated to UMRs and LMRs:** (a) UpSet plot representing the overlap with functional elements and LMRs. Only functional elements that overlap with any LMR are included in the figure. (b) Mutation estimates in functional element free UMRs for DNA repair deficient tissues. (c) Methylation level of all UMRs and others that do not show overlap with promoters (proless) or loop anchors (lapless) or both (lapproless). (d, e) Methylation level of all promoters and loop anchors, overlapping and not overlapping UMRs and other functional elements for LAPs. Methylation level is represented as the median average value of all cases. Shadow represents the 95% binomial confidence interval of the median. (f) Mutation rate estimates in promoters that significantly overlap with a UMR (> 200bp) and all promoters. (g) Same as in f but for selected tissues and stratifying the promoters according to the expression bins.

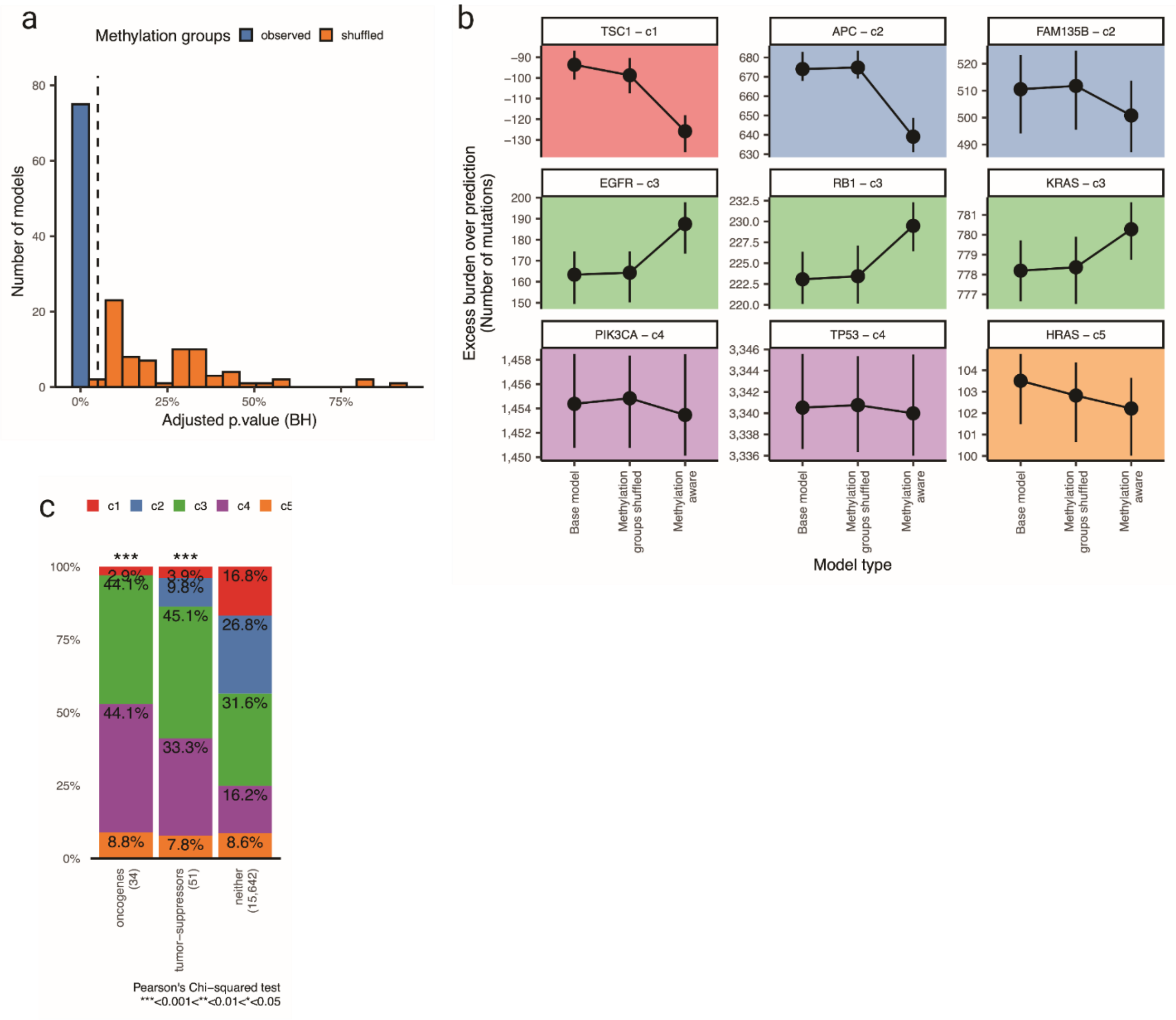


**Extended Data Fig. 5. Gene mutation burden prediction with methylation aware gene classes:** (a) Significance measured by log-ratio test (see Methods) of a model including the observed or a shuffled version of the methylation aware gene groups. Vertical dashed line indicates significance threshold (0.01). 6 out of 75 models with shuffled groups showed significance. (b) Selected examples of burden excess, mutations observed versus mutations expected, for some cancer driver genes. Colors of the box represent their gene group. Note axis are different in each box.
