## Supplementary Methods for "Mutation rate heterogeneity at the sub-gene scale due to local DNA hypomethylation"

### Supplementary Methods for Mas-Ponte and Supek (2023)

*Mutational signature assignment*

The downloaded somatic mutations from whole-genome sequences (see Methods) were tallied and classified according to their trinucleotide context and their alternative base. COSMIC signature profiles and tissue exposures (V3.3) we downloaded directly from the cosmic website at (cancer.sanger.ac.uk/signatures/sbs/). A mutational signature was assigned to a tissue if at least 1 sample in the cosmic dataset contained that signature. Of note, some of the samples in the cosmic signatures dataset are also included in our set, but their direct exposures were not used from this file. Signature 1 and 5 were included in the assigment to all tissues. MSI and POLE deficient samples were treated independently of their tissues of origins and signatures associated with their phenotype were included, in brief, for MSI samples we included signatures 6, 15, 21, 26, 44, 14, and 20; and for POLE deficient samples we included 10 (a, b, c and d), 14 and 20. For each tissue, the matrix with the mutational profile of each sample was computed and fitted to the assigned cosmic signatures using SigLasso[1] which implements a lasso regression mutational signature fitting that forces sparsity. Results from the SigLasso fitting were then used as exposures for the rest of the analysis.

For every sample in our dataset, we used the signature exposures obtained from SigLasso fitting to obtain the probability of a given mutation to be caused by a given mutational signature. In brief, the exposure in each sample was split to the 96 mutation categories according to the original mutational profile (weight of every trinucleotide subtype) and each feature was normalized within every sample so that every mutation class had a given probability to be associated to any of the used mutational signatures. Thus, using this approach, we could estimate the probability of a mutation subtype to be associated with a given signature. If the signature was not present in a sample the probability was then zero.

Then, to classify the raw mutation calls, we used these probabilities to sample a unique signature to each mutation according to the probability assigned to its mutation subtype. This process allows us to classify raw mutation calls to distinct mutational signatures and allows us to pool mutations generated by the same process across different samples and different tissues.

*Gene models and average expression values*

Gene models for assembly GRCh37 were downloaded from the GENCODE releases website (https://www.gencodegenes.org/human/) for version 19. For each gene, a single transcript was selected according to the TREGT gene list that uses a combination of CDS gene length and expression level to select the most appropriate isoform for each gene. The list is available in (tregt.ibms.sinica.edu.tw) and in ref[2]. Because the annotation reference of this resource was based on gene models from the GENCODE version 26 the gene ids and transcript ids that did not match with our annotation were discarded, leaving a total of 15,715 genes (Supp. Table 3). Genomic coordinates for the TSS (5' end), TES (3' end) and gene length were derived directly from the v19 GENCODE annotation file.

Transcription levels for all genes were downloaded directly from GTEX website (version V8, GTEx_Analysis_2017-06-05_v8_RSEMv1.3.0_transcript_tpm.gct.gz) in TPMs and averaged globally for all available tissues within the dataset. This analysis then yielded one average expression value per gene that was binned in three equal size terciles.

*Mutation rates estimation using Negative Binomial regression*

Throughout the analysis in this study, the estimation of the local mutation rate of a specific region of interest (ROI) was performed using a Negative Binomial regression. This includes analysis on the local mutation gradients (Fig. 1), and hypomethylated regions (Fig. 3 and 4).

In brief, mutations were stratified according to their trinucleotide content and their overlap with a specific region. Then, mutations were tallied over those features effectively pooling across types of regions. Likewise, trinucleotides of the reference sequence were also tallied in the ROIs to determine the nucleotides at risk for each context. These values were used as an offset in the regression allowing control for the sequence context in the regression.

The function MASS::nb.glm is then used to perform the negative binomial regression over the data table. The total number of rows is equal to the number of contexts used (96) multiplied by the region channels. This step leads to a formula such as:

$$\mathrm{M} \sim \mathrm{R} + \mathrm{offset}(\ln(\mathrm{N}))$$

Where M is the number of mutations, R represents the region of interest and the N represents the nucleotides at risk corresponding to each trinucleotide.

To calculate the mutation rate gradients, each bin within the genes represent one region of interest and the resulting mutation rate is measured relative to a reference window located 2 units of the outward expansion of the bins (4kb) downstream of the TES. The reported mutation rates are thus all relative to this bin. To measure the mutation rate at UMR or LMRs we compared the mutation accumulation at the region of interest (ROI) against their flanks. We defined flanks as the regions separated from the ROI by 1 width. Each flank had half of the width of the original ROI. This essentially translates to splitting the UMR/LMR in two halves and moving each section one width in the corresponding direction. Mutation rates are always represented as the ROI over flanks.

Using these designs for both gene gradients and unmethylated regions, the reference region and the ROIs will likely be located in the same replication time domain minimizing the need to control for this co-factor. At the same time, separating these regions by a factor higher or equal than one width allows us to detect clean signals which can not be underestimated due to loose ends when segmenting the genome.

The resulting estimate from the regression is given as the natural logarithm of the odds ratio which can then be later transformed to logarithm in base 2 or as a percentage change. If not stated otherwise, mutation rate enrichments on figures are displayed as a base 2 logarithm.

*Methylation data sources*

To maximize the genomic coverage of the DNA methylation data, we gathered whole genome bisulphite sequencing (WGBS) from publicly available datasets, in brief, the Roadmap epigenome project (see https://egg2.wustl.edu/roadmap/web_portal/ ) and the ENCODE data portal (see https://www.encodeproject.org/).

Data from the Roadmap project consisted in all sets with available WGBS data. They can be accessed in Supp. Table 2. Downloaded data consisted in fractional methylation data ( FractionalMethylation ) which contains information about the methylation of each sufficiently covered CpG in a percentage value. We also downloaded files containing genomic coverage of each CpG.

Similarly, all WGBS available data from ENCODE were downloaded. All files were in bedMethyl format derived from the output of Bismark[3] in the ENCODE main processing pipeline. This format also contains the methylation levels of all sufficiently covered CpG in a percentage. In addition, the same format also contains information about the coverage of each CpG dinucleotide. Accession codes from these files are available in table. The ENCODE datasets were only available in the hg38 assembly and were translated to hg19 (using liftOver) to match the rest of the analysis. LiftOver statistics can also be found in Supp. Table 2.

Whole genome bisulfite datasets that were excluded in the UMR analysis (see below and Supp. Fig. 6 and 7), were also excluded in this analysis.

##### *Clustering of methylation profiles in gene bodies*

From the downloaded WGBS datasets, the average methylation value for every available CpG dinucleotide was computed within tissue groups (brain, blood and solid). The solid average values were used for this analysis.

Gene bodies were extracted from TSS to TES , thus including 3’ UTRs, coding sequences, introns and 5’ UTRs. For each gene body, the analyzed regions were located around either ends. These ends were expanded 3kb outward, upstream for the TSS and downstream for the TES, and 5kb inward, in reverse order. These sections were divided in 50bp sections. If genes were shorter than 5kb, the bins were further expanded from each direction. Methylation averages were then extracted from each bin using the calculateMatrix tool in deeptools[4] generating a matrix with TSS and TES concatenated bins as columns and genes as rows.

The resulting matrix was factorized using a PCA (from the R FactoMineR package) with no scaling. The NA values in the matrix, representing bins with no methylation signal, were imputed automatically using the mean value of the column. Significance for the number of principal components was extracted comparing to a broken stick model (from the R vegan package), which simulates a non-signal scenario. The resulting coordinates of each gene for the top three principal components were grouped using medoids clustering (function cluster::pam in R). The number of clusters selected (k = 5) was chosen from a range (2 to 7) after visual inspection of the resulting methylation profiles and genomic characterization (Supp. Table 3). Although a selection process based on silhouette index and sum squared of the residuals was also performed, the continuous characteristics of the clustering and the lack of defined numerical limits made this approach too conservative. The reader might interpret these clusters as groups of similarly methylated genes with similar genomic properties.

To extract the methylation profile of every gene cluster, genes were grouped according to their assigned cluster and the average value was computed for each bin. This profile is indicative of the different methylation profiles in each group. Meta profiles of the methylation along the gene body were computed using the computeMatrix utility from deeptools in reference point mode. The average methylation profiles for each gene group were performed using *in house* scripts which also included the measure of a confidence interval. The confidence interval of the median is measured using the indices of a binomial distribution with the given sample size equal to the number of rows tested, here, the number of genes in a specific cluster. Confidence interval levels are always 95% two-tailed if not stated otherwise.

For the genomic characterization of the profiles, genes were tested for local enrichment of histone marks, promoters, and enhancers and chromatin states (see below). Promoters and enhancers were downloaded from the FANTOM[5] V5 dataset but pooled across all expression levels for this analysis.

##### *Chromatin states, CpGi and HGC association*

ChromHMM states were downloaded as a bed file from the Roadmap data portal at egg2.wustl.eduroadmap/data/byFileType/chromhmmSegmentations/ChmmModels/core_K27ac/jointModel. The core_K27ac model was selected for sample E017, that corresponds for IMR90 cell line, and used throughout all the analysis.

When measuring enrichment of chromatin states in methylation aware gene groups, relative Fig. 2F, a matrix encompassing the different chromatin states and different groups was build with the size of the overlaps in basepairs. A $\chi^2$ test (chisq.test) was performed on the resulting matrix that worked as contingency table. The values represented in the figure represent the observed overlap over the expected from this test. In order to asses the significance of a specific set of chromatin state and gene group, a partitioned $\chi^2$ test using a 2x2 contingency table that included the selected gene group and chromatin state and was compared among all other.

Analysis of the overlaps of these groups against CpG islands and high GC content genes were performed likewise.

The division of genes categories according to the CpG content in their promoters was extracted from the supplementary material of ref[6] for CpGi genes and was calculated as in ref[7] for the HCG genes. In brief, CpG instances were tallied in each promoter and normalized against its CG content. This measure was then modeled by a Gaussian mixture model (using mclust package) with two components.

##### *Histone mark ChIP-seq datasets*

Tissue specific data for the selected tissue groups (solid, blood and brain) were downloaded from the ENCODE main data portal (https://www.encodeproject.org/). From each of the selected groups of tissues we obtained 3 reference experiments (reference epigenomes). If available, data from primary tissues was obtained. If not available, data from cell lines and primary cell cultures was used.

We obtained a total of six histone mark signal for every experiment: (i) H3K4me3 for TSS and promoters; (ii) H3K4me1 for enhancers; (iii) H3K27ac for active promoters and enhancers; (iv) H3K9me3 for heterochromatin; (v) H3K36me3 for gene bodies of expressed genes and (vi) H3K27me3 for bivalent transcription and Polycomb marked genes. The signal obtained measured fold change over control which is equivalent to the chip-seq signal value over the input in the experiment.

For the 3 samples included in each group, we averaged the signal using UCSC tools ( bigWigMerge). We then combined the averaged signals with the different gene groups to obtain a metaprofile using computeMatrix from the deeptools[4].

##### *Unmethylated and partially unmethylated regions*

In order to call significant unmethylated regions (UMR) we used *MethylSeekR* from bioconducor[8] implementing the default processing workflow suggested by the authors in the vignette. In brief, SNP positions are first removed from the set (see Supp. Table 2). PMDs were detected by using the shortest chromosome with at least 150 probes as a training set. CpG islands were downloaded from UCSC table query. These datasets were then used to calculate the FDRs for the detected UMR segments. A threshold of 4 CpG positions in each segment and at least a smaller than 50% methylation value was required. If the FDR value at these conditions was lower than 5%, the samples were automatically discarded. If the total number of CpG islands considered was smaller than 25M the samples were also discarded. Non-autosomal chromosomes were removed (Supp. Table 2).

This process was run for every sample in our dataset individually. UMRs extracted from each set were then translated in a matrix format, containing a binary encoding (1 or 0) if a specific locus was included or not in that sample. This matrix was factorized using tSNE (from the Rtsne package) with 25 perplexity. The resulting grouping was inspected for biological coherence, samples that were not grouped with its tissue group were manually excluded for further analysis (see Supp. Table 2).

For each tissue group (solid, brain, and blood), individually detected UMRs were pooled into a union set which contained all UMR loci from every experiment and then reduced to avoid overlaps. If not stated otherwise, these are the sets used for all analysis when compared to mutation calls. A full union set was also generated from the union of all sets together. Each union set for every tissue was then used to compare with the other tissue groups and the UMRs which were specific to that tissue group, not present in others, were selected as tissue-specific.

UMRs and LMRs from other studies were also downloaded to be used as reference sets in this analysis. From ref[8], the downloaded datasets were in the supplementary material and were pooled for both UMR and LMR classes. These experiments included mostly cell lines from blood tissues, reprogrammed cells, adipose tissue and fibroblasts. This dataset was originally downloaded in hg18 and then translated into hg19 with liftOver. Of note, software used to call UMRs in these datasets was the same as the one used for the downloaded WGBS data. Data from ref[9] was also downloaded from the supplementary material and pooled across different available categories. The UMRs were divided into Canyons, cUMR (conserved UMRs) and either healthy or tumor specific UMRs. If not stated otherwise, the conserved UMR dataset was used for all the analysis in this study. The distinction of UMR versus LMR was not available in this study and thus was not included in our analysis.

##### *Functional elements intersection with UMRs and LMRs*

Enhancer data based on CAGE data was obtained from the FANTOM dataset[5], version V5 (https://fantom.gsc.riken.jp/5/datafiles/latest/extra/Enhancers). They were posteriorly divided into terciles using the predefined categories in the downloaded data, with t3 indicating a higher expression level (in TPMs) and t1 indicating the lowest. As in ref[9], superenhancers were downloaded from the supplementary material in ref[10]. From the available sets we used primarily the superenhancer track marked in red. The UCSC gene model, available in the bioconductor package TxDb.Hsapiens.UCSC.hg19.knownGene , was used to define promoters and the 5' genic sections. Promoters were defined as the 2kbp upstream of the TSS with no upstream segment, and the 5' genic sections were defined as the 2kbp downstream of the TSS.

These functional elements were compared against different sets of UMRs for three different sources of methylation data: from ref[8,9] and the set gathered in this study. The enrichment measurement is based on a fisher exact test of the overlapping bp between 2 types of regions. Thus, if a feature is less specifically overlapped against another, the odds' ratio will decrease even if many nucleotides of the sparser one are covered.

##### *Tissue specific analysis of the mutation rate*

UMRs and LMRs were also classified in three tissue specific sets, blood, brain and digestive. 3 sets for each group were used in the comparison of sites. UMRs that were present exclusively in one, two or three datasets within the same group were considered as tissue specific. Regions that were present in all the datasets included in the analysis were considered as consistent and the regions that did not meet any of these criteria were labeled as other. Then, mutation rate was computed as above by matching the methylation based UMRs with samples from brain, myeloid blood (to avoid introducing the lymphocytic samples) and colon cancers to all tissue specific UMR/LMR(s).

##### *Measuring selection through burden prediction*

To estimate selection in genes we used whole exome data available from the mc3 dataset[11] from TCGA. A total of 10,295 samples were downloaded from (https://gdc.cancer.gov/about-data/publications/mc3-2017). Only single-nucleotide variants in autosomes were kept for further analysis. Mutations in these sites were assigned to genes from the GENCODE v19 annotation as in other analysis. We downloaded the epigenomic-based principal components from the dNdScv[12] method to use as a covariate of a base model with no methylation information from (https://github.com/im3sanger/dndscv/blob/master/data/covariates_hg19.rda?raw=true). We then compiled the coding sequence of each gene and tallied their trinucleotide sequence contexts and normalized them per gene. The total width of the gene was also compiled.

A negative binomial regression was designed to predict the total burden of the gene, defined as the number of mutations accumulated in that gene in a pan-cancer setting. The implementation of the negative binomial in the MASS package was used as above. Predictor features included in the base model were the percent of each trinucleotide in a gene and the epigenetic predictors downloaded from dNdScv. The model also used the total size of the gene as an offset of the regression. Two additional methods were tested, one containing the methylation aware groups of genes defined above (Fig. 2, Supp. Table 3) from the DNA methylation profiles of genes and another one containing a shuffled (resampled without replacement) version of the grouping. The shuffled model was used as a negative control across the analysis as it maintains the same degrees of freedom and complexity as the original one.

Genes associated with cancer development (henceforth driver genes) were downloaded from the mutpanning[13] website (http://cancer-genes.org/). Only genes with a significant association in at least 2 cancer types were considered. All genes in this list were considered as positive set and thus were excluded from the train/test split. To train the model, we repeated a 5-fold cross validation 15 times yielding a total of 75 iterations or models. Prediction was then performed for the set of genes either in the testing set or the positive set and the excess of somatic mutations was then calculated as the expected number of mutations minus the observed. Thus, a higher excess implies a higher burden than expected and thus plausibly positive selection.

In order to measure significance of a single feature of the model we used the fitted version and calculated a sequential analysis of deviance table from the anova.negbin function in the MASS package. This function performs a log-likelihood test for the deviances of the model with and without every feature. In our analysis we extracted the significant values from the methylation aware gene groups and its shuffled form only and then corrected them for multiple testing using Benjamini-Hochberg.
