## Supplementary Figure Legends for "Mutation rate heterogeneity at the sub-gene scale due to local DNA hypomethylation"

**Supplementary Figure Legends for Mas-Ponte and Supek (2023)**

Supplementary figures are available as external PDF files.

**Supplementary Fig. 1. Gene gradient plots for individual regressions.** Coefficients (in points) and 95% CI (as vertical lines) of the regression values from a gene gradient analysis. Each point represents a 250 bp long bin and are distributed along the gene body in a region around the 5' end, the central location and the 3' end (same as in Fig. 1). Values represent the coefficients in a pan-cancer setting, thus, all mutations assigned to a given signature across cancer types are considered. Furthermore, genes are split across average GTEX expression level in equal sized terciles. Eq1 represents lower expression levels and Eq3 higher. (Relative to Fig.1A)

**Supplementary Fig. 2. Coordinates of individual points in the gene-gradient PCA.** Coordinate values of every row included in the PCA of the gene gradients. Mutations were stratified in tissue, expression level, and signature. Thus, every square contains information about a single instance in the PCA. Empty squares result from no mutations assigned in that tissue of origin and signature, or alternatively, due to a failed regression because of low counts. (Relative to Fig.1C and Extended Data Fig. 1E)

**Supplementary Fig. 3. Contributions of each instance into the gradient PCA analysis.** The contribution percentage of every instance in the PCA to the five top components. Contributions are represented as percentage so negative and positive coordinates can have both higher contributions although opposite trends. The total sum adds up to 100% for each component (Relative to Fig.1C and Extended Data Fig. 1E)

**Supplementary Fig. 4. Histogram of average methylation across gene groups** (a), further subdivided in distinct gene sections (b), the 3' and 5' UTR exons and the first, second and third coding exons from the 5' end of the gene body. (c) Methylation average values of genes encompassed in the mid expression and mid size category subdivided by introns and exons. Dashed sections represent extended 200bp from the section end.

**Supplementary Fig. 5. Gene gradient mutation rate coefficients for methylation aware gene groups.** Mutation rate gene gradients (as in Fig. 1) were calculated for each mutational signature in groups of genes divided according to their methylation profiles (from Fig. 2). The heatmap shows the coefficient values (in ln scale) of the 5' end, central and 3' end gene sections. Rows are clustered and annotated with the methylation group of genes where the regression was performed.

**Supplementary Fig. 6. Excluded WGBS samples from MethylSeeker output**. MethylSeeker FDR for UMR/LMR calling at predefined cutoffs (solid lines, see Methods). Each row contains a selected experiment. (top) Global false discovery rate for a single sample. (bottom) Total number of CpG fragments obtained from every sample. (Relative to Methods and Supplementary Table 2).

**Supplementary Fig. 7. Manually excluded WGBS samples.** tSNE plot from UMR/LMR detected loci in WGBS samples (see Online Methods). All samples showed had passed the previous trhresholds in MethylSeeker. After manual inspection, samples that did not cluster with the intended tissue were manually excluded (marked with a cross) and removed from the analysis.
