## Supplementary figures and images for "Mutation rate heterogeneity at the sub-gene scale due to local DNA hypomethylation"

### Supplementary Figure 1

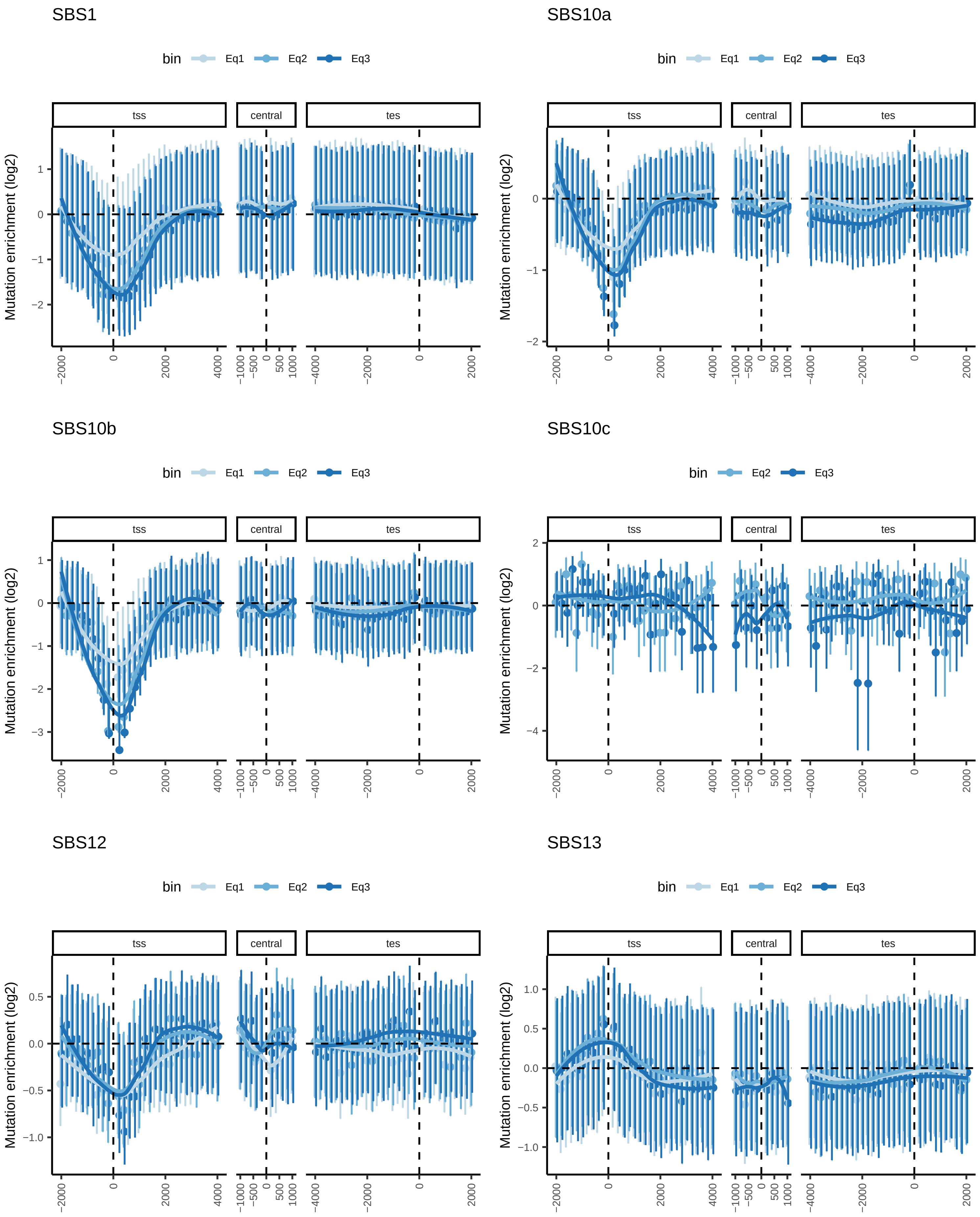

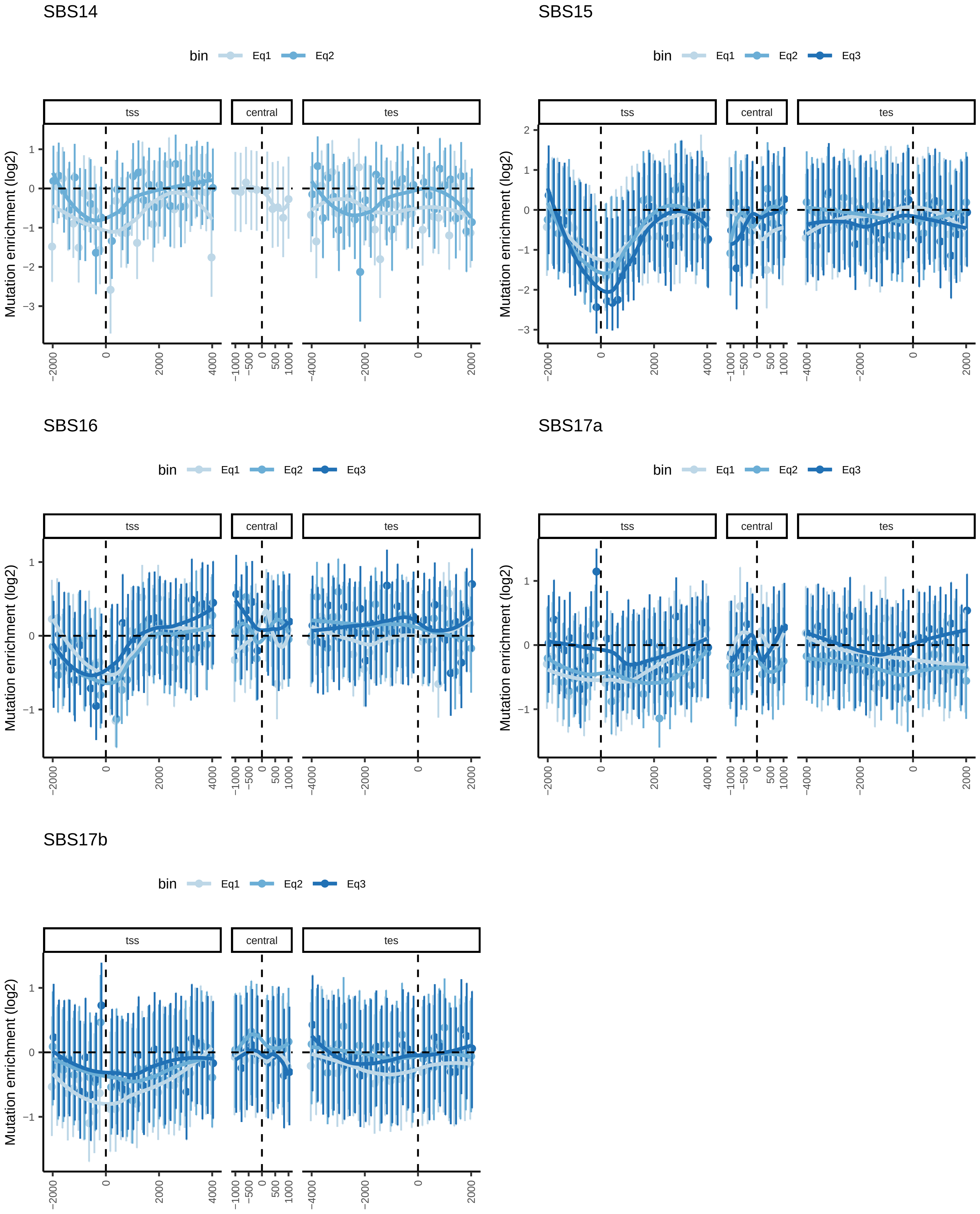

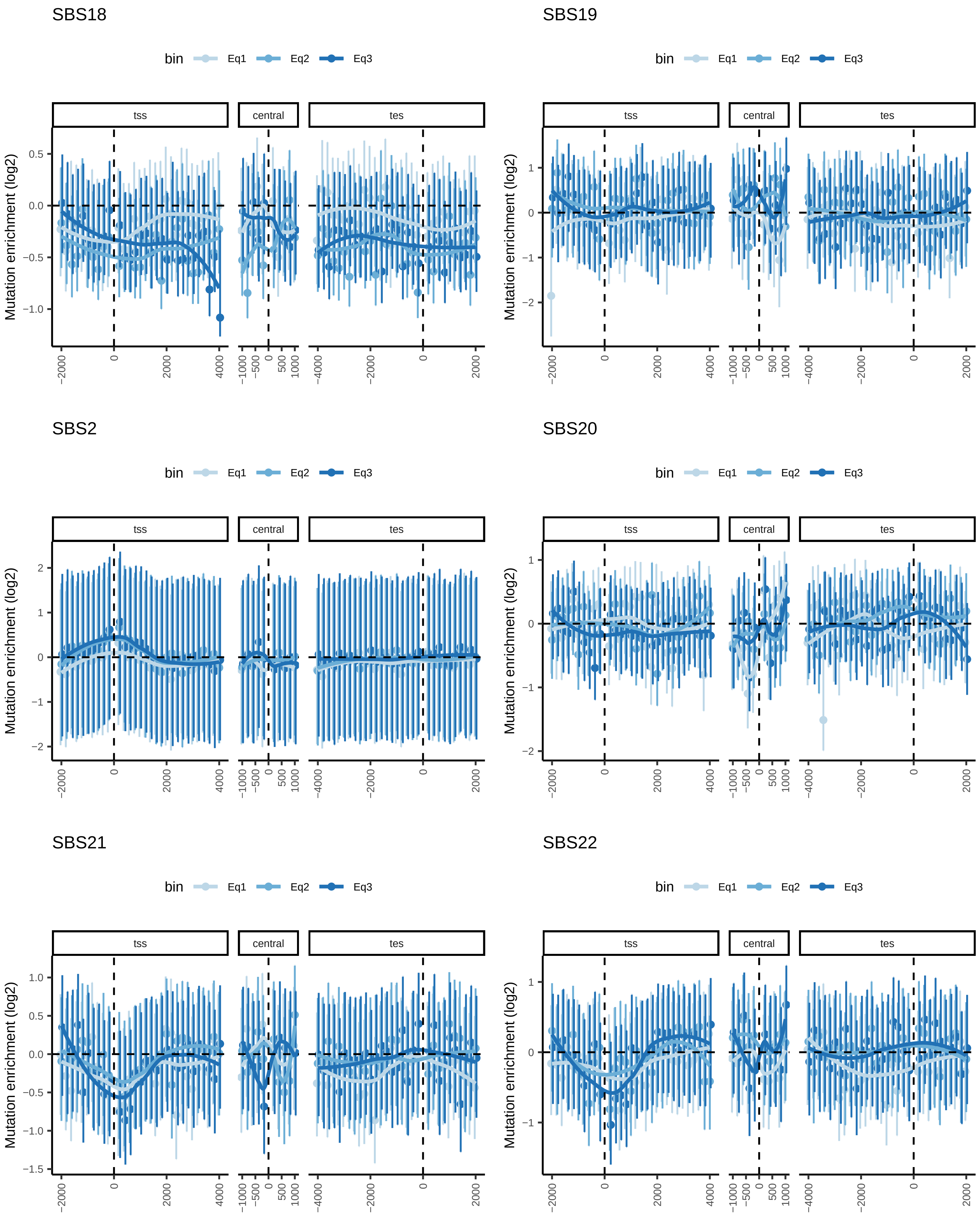

SBS24

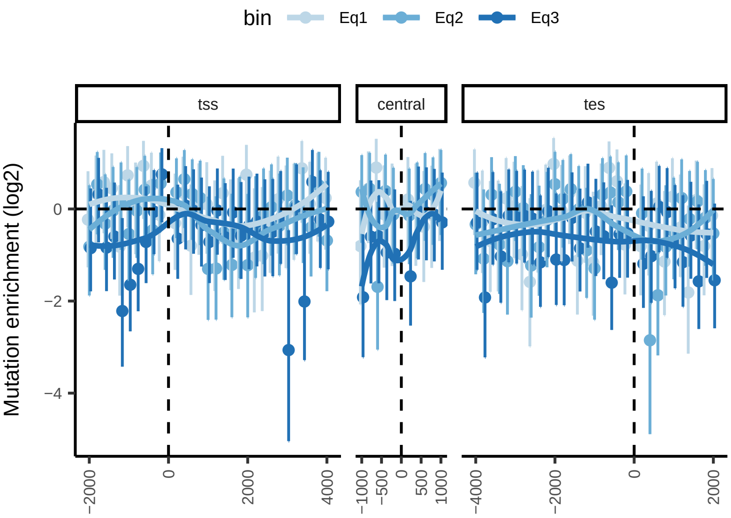


SBS25

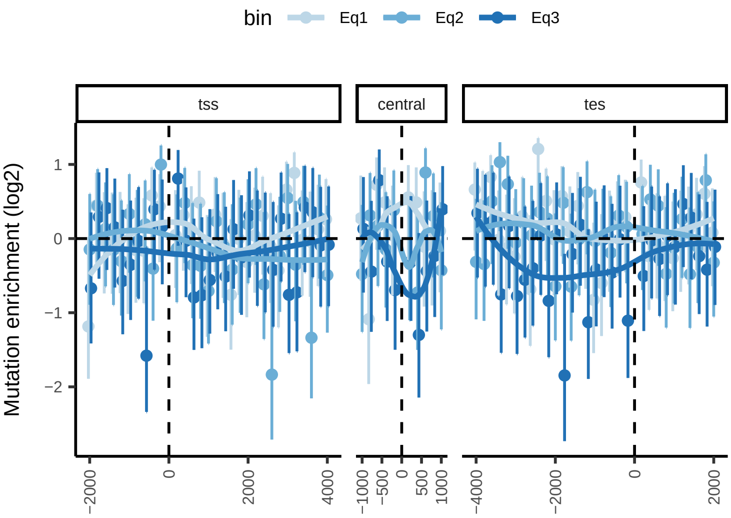


SBS26

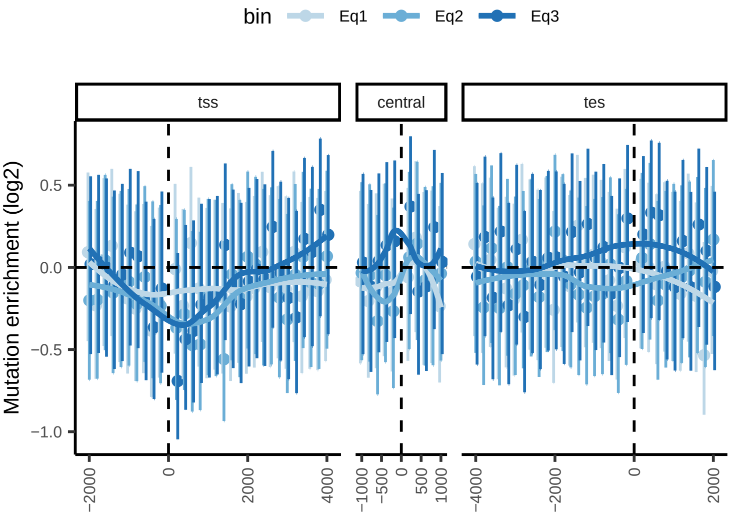


SBS28

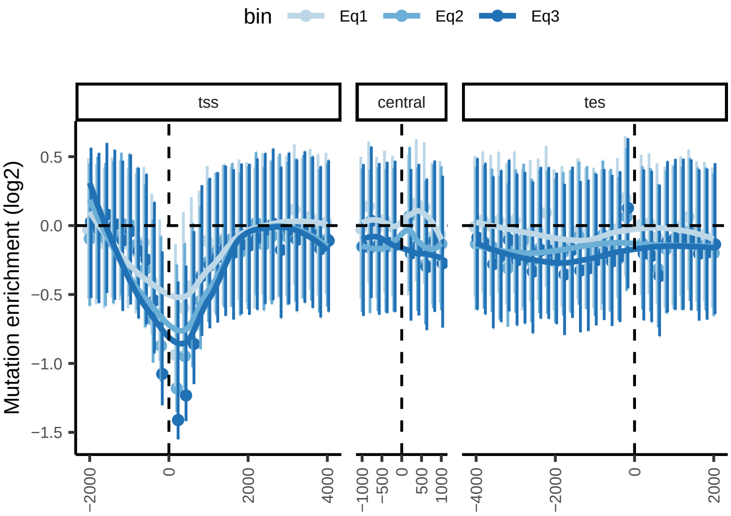


SBS3

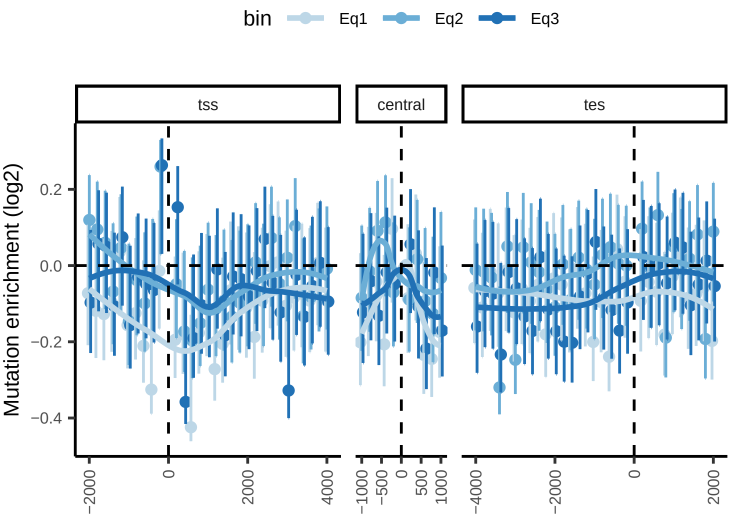

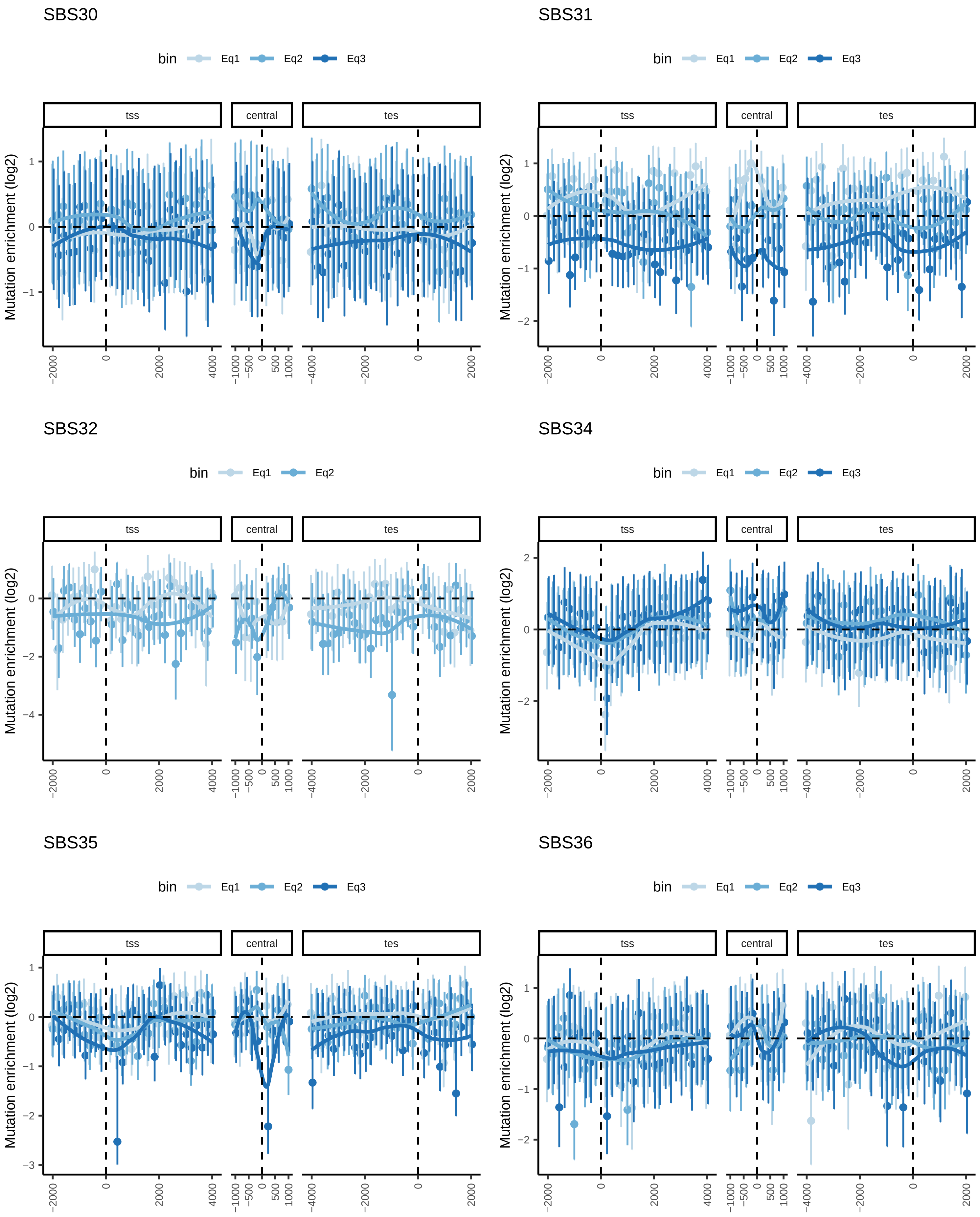

SBS37

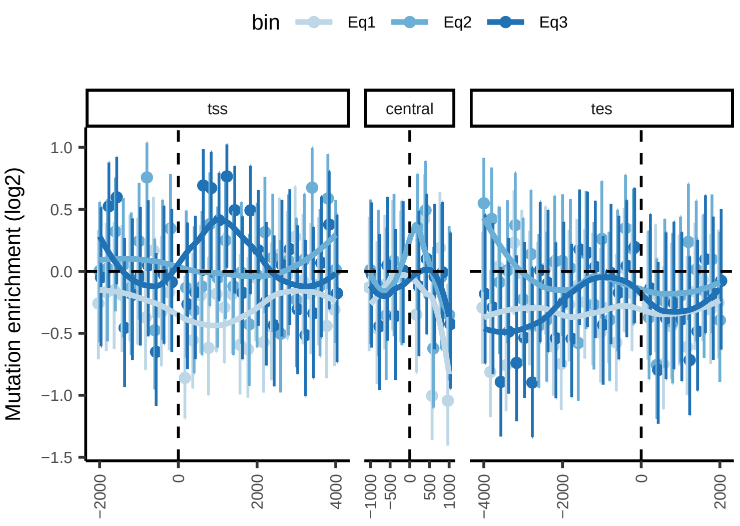


SBS38

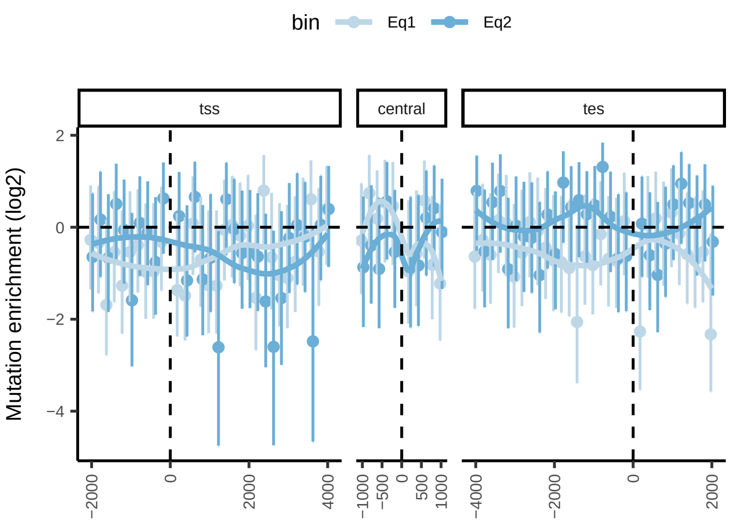


SBS39

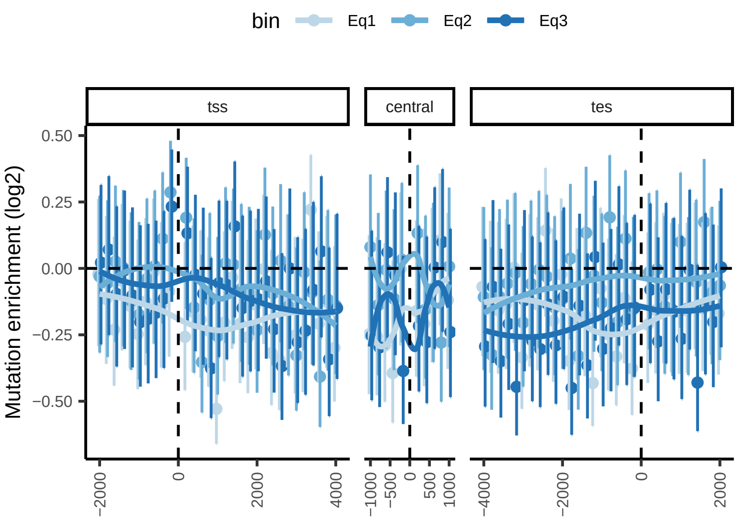


SBS4

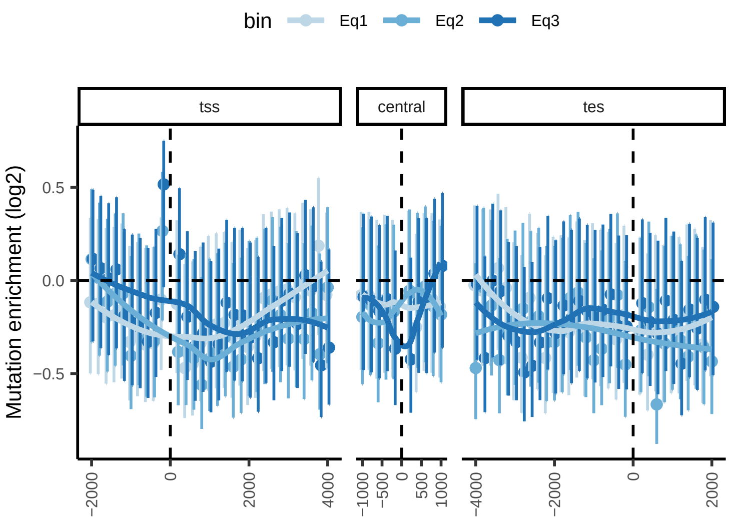


SBS40

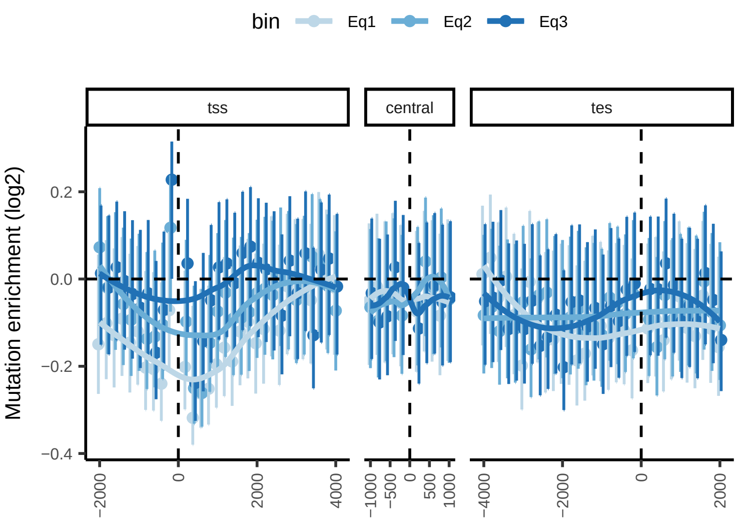

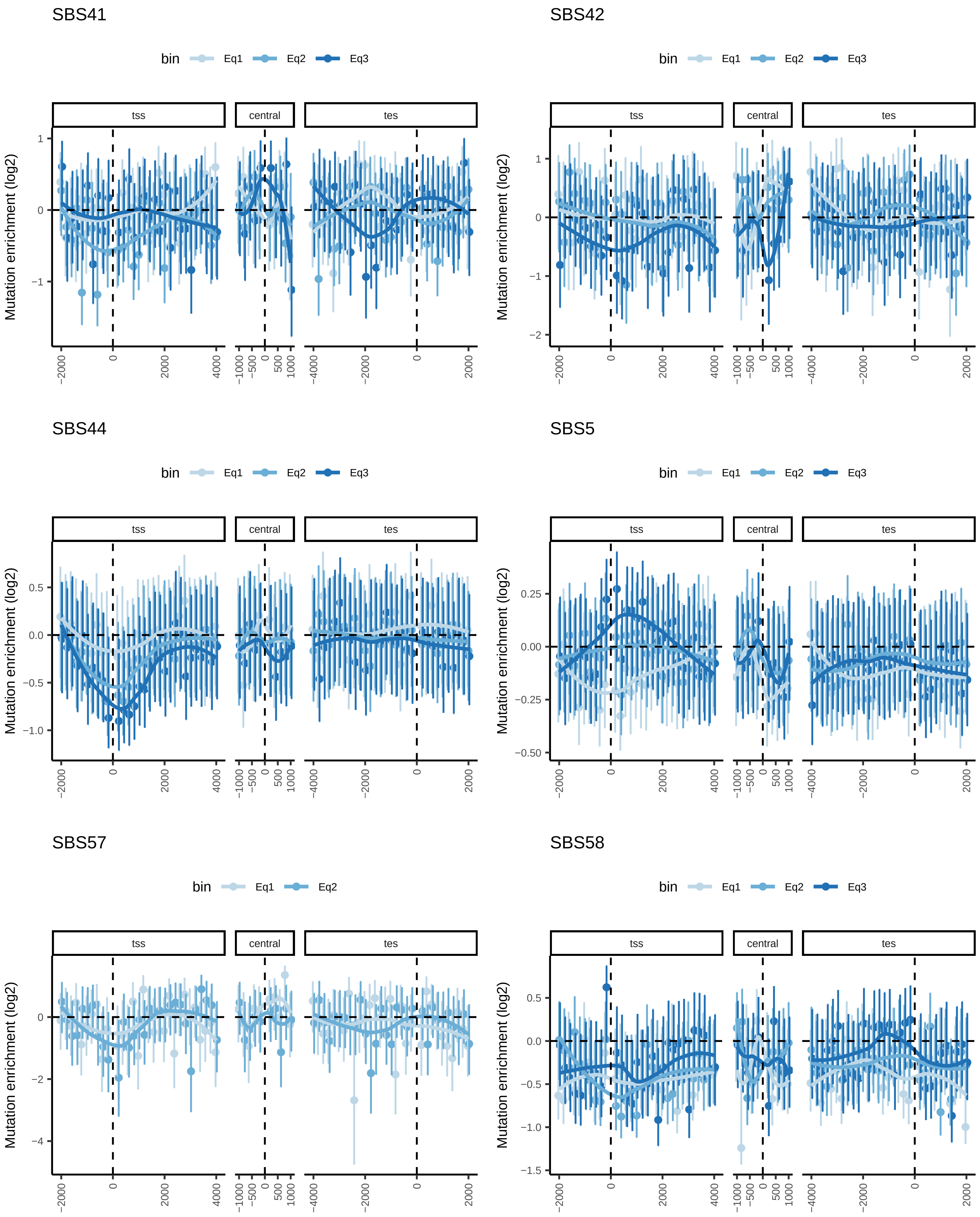

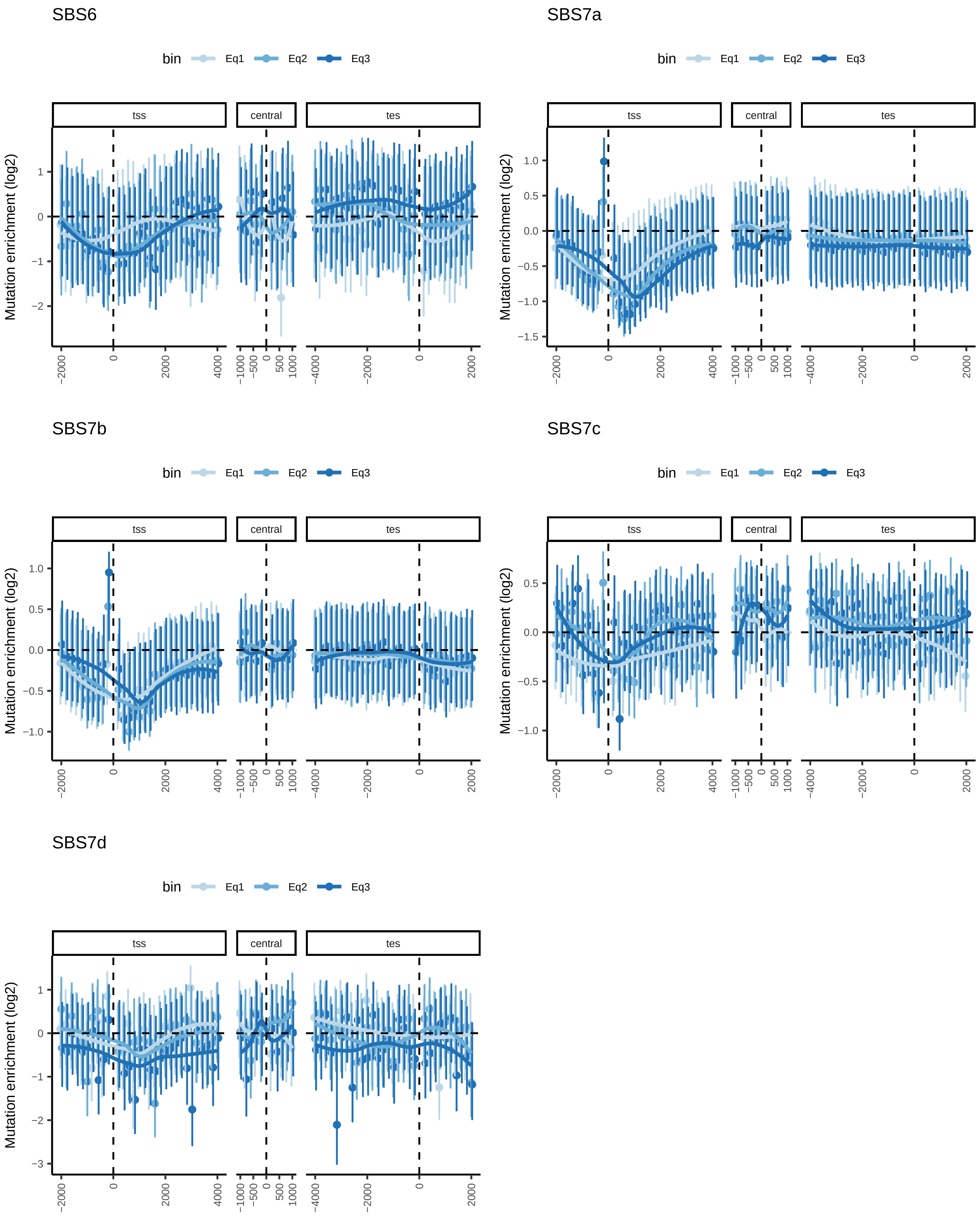

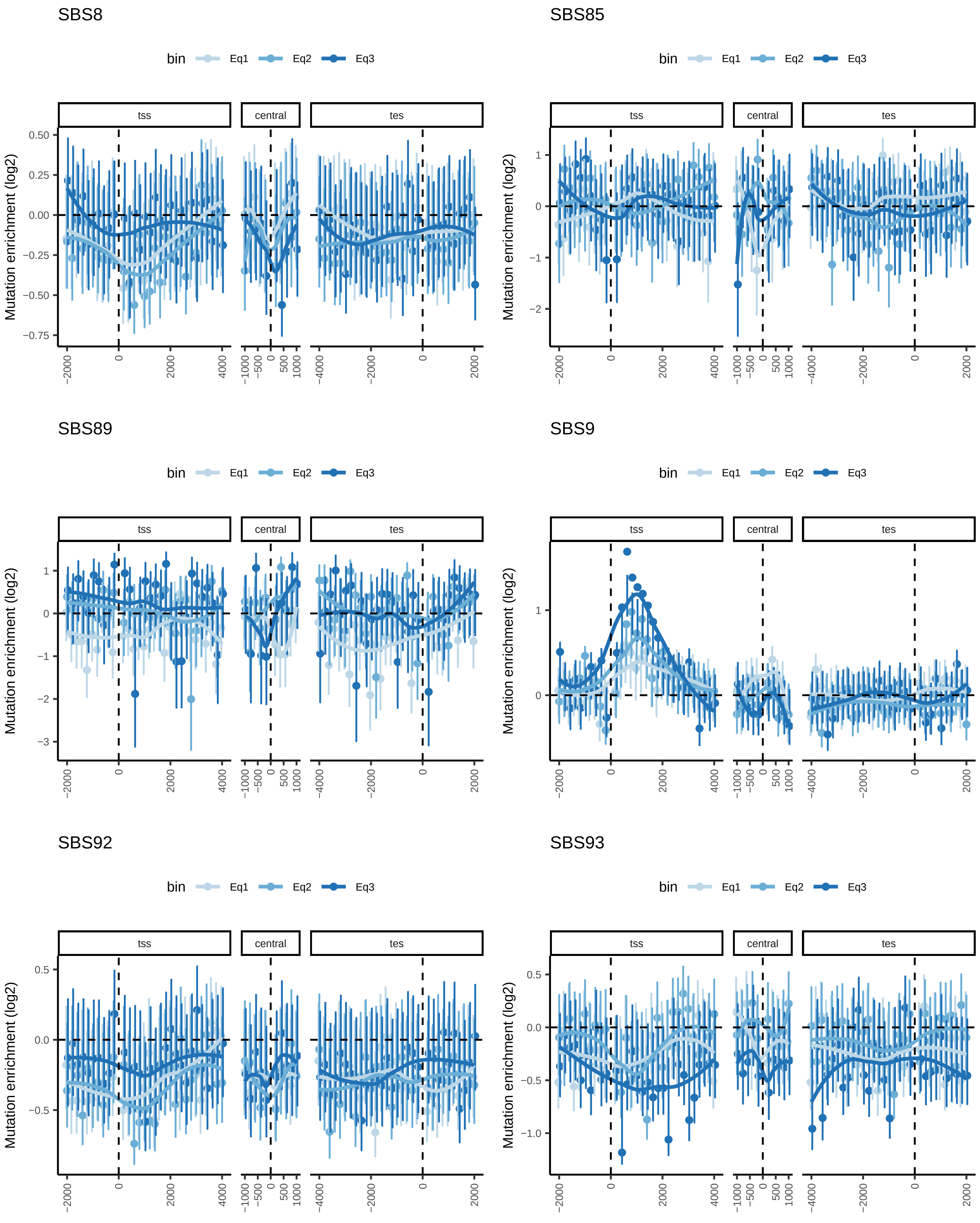

### Supplementary Figure 2

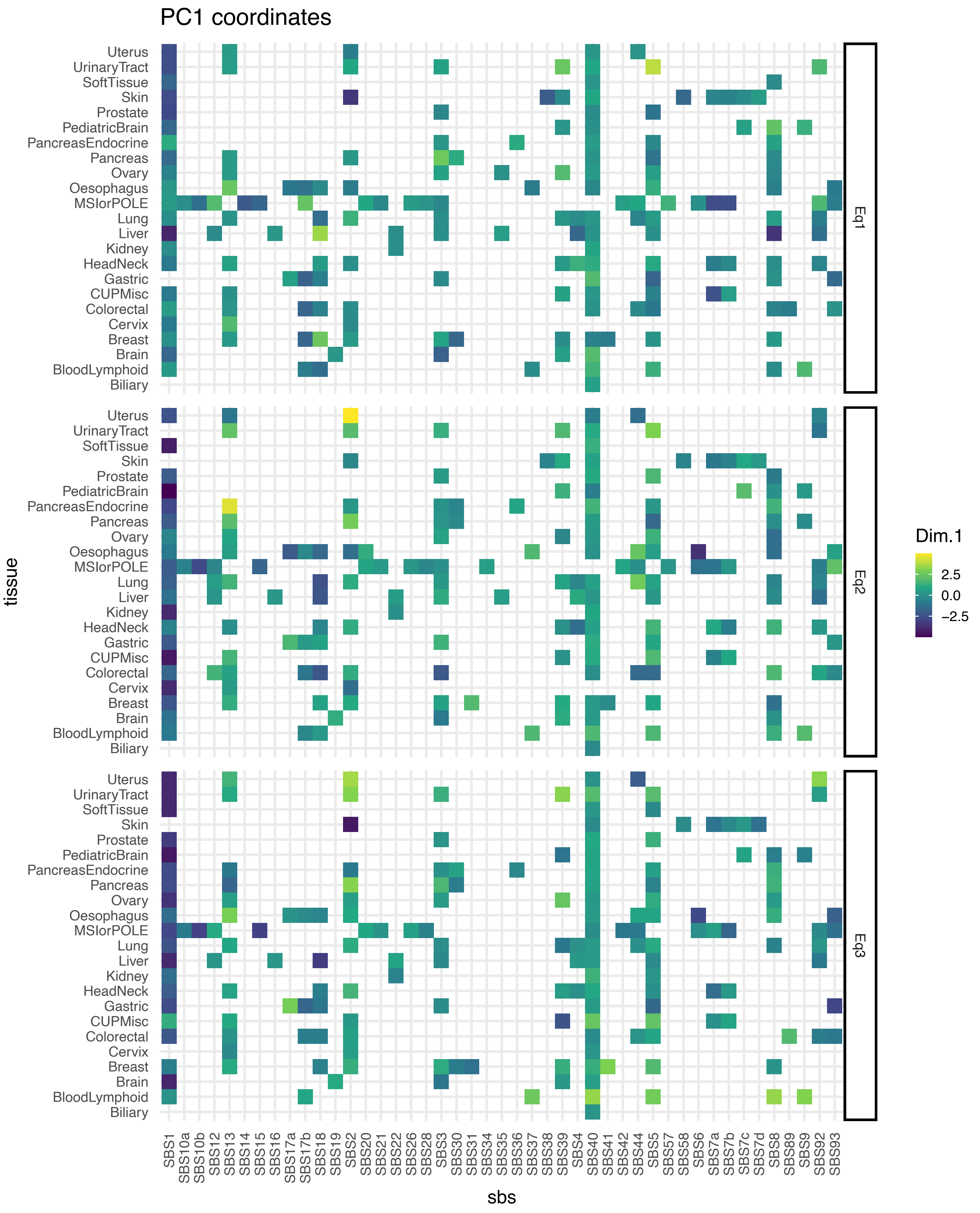

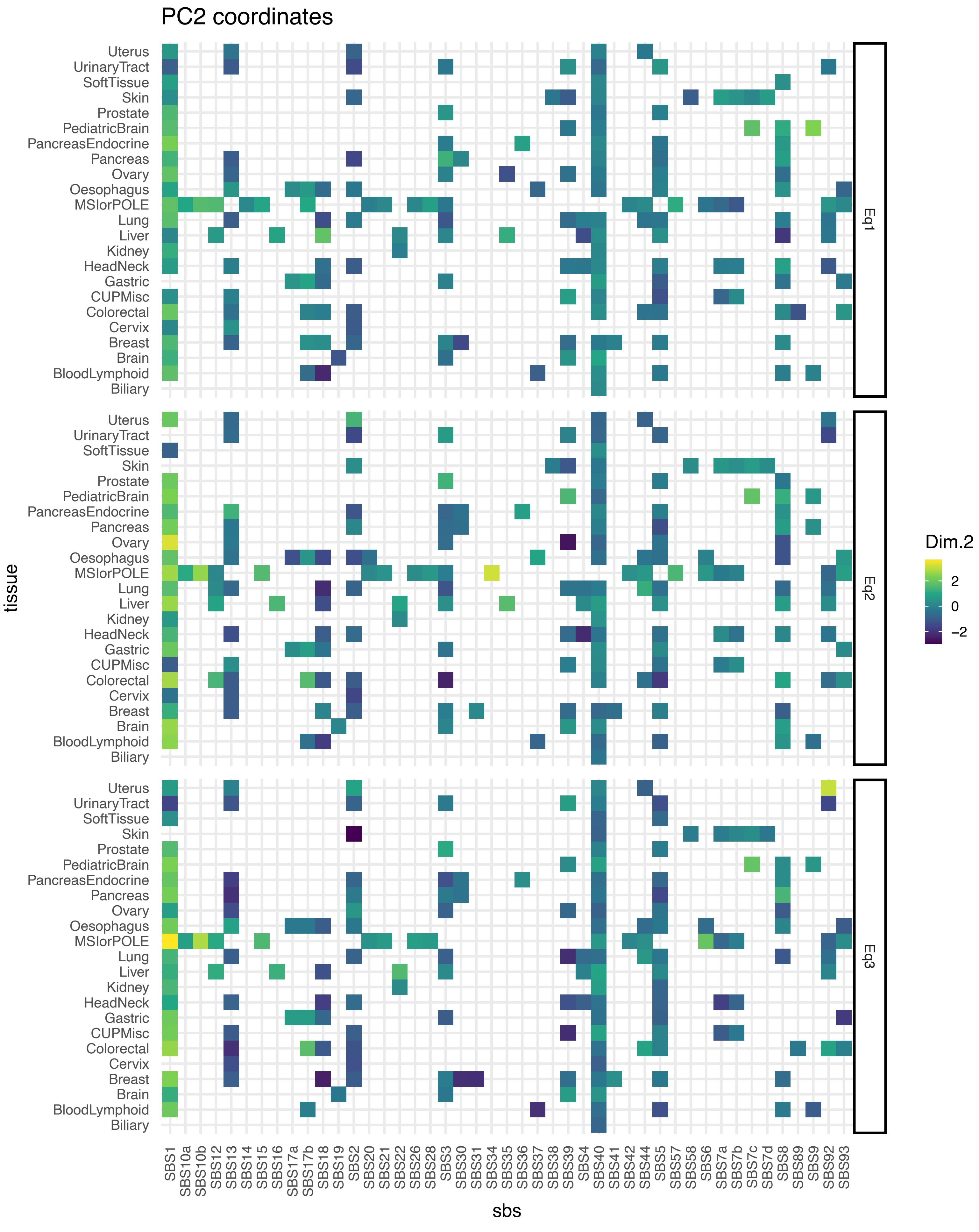

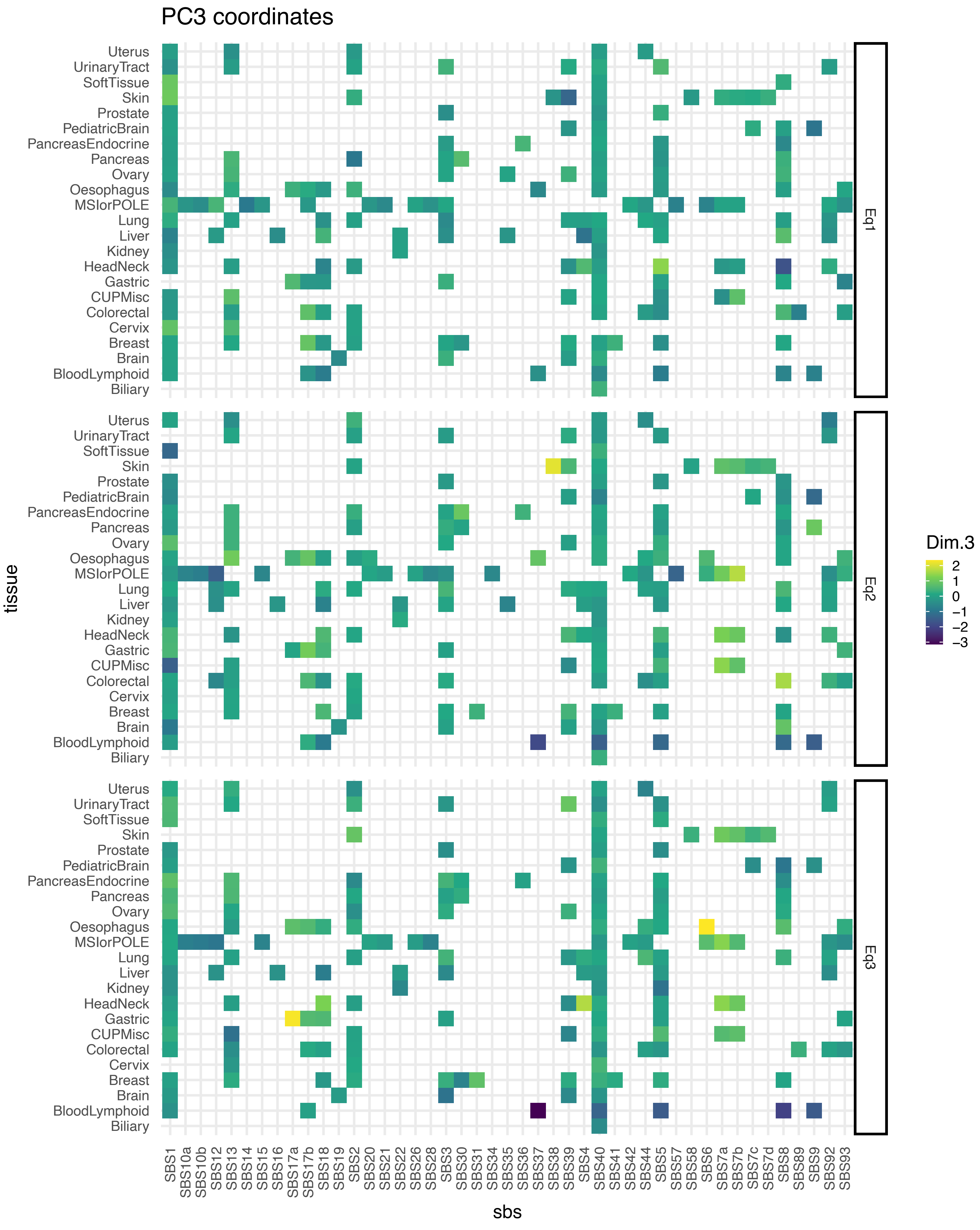

### Supplementary Figure 3

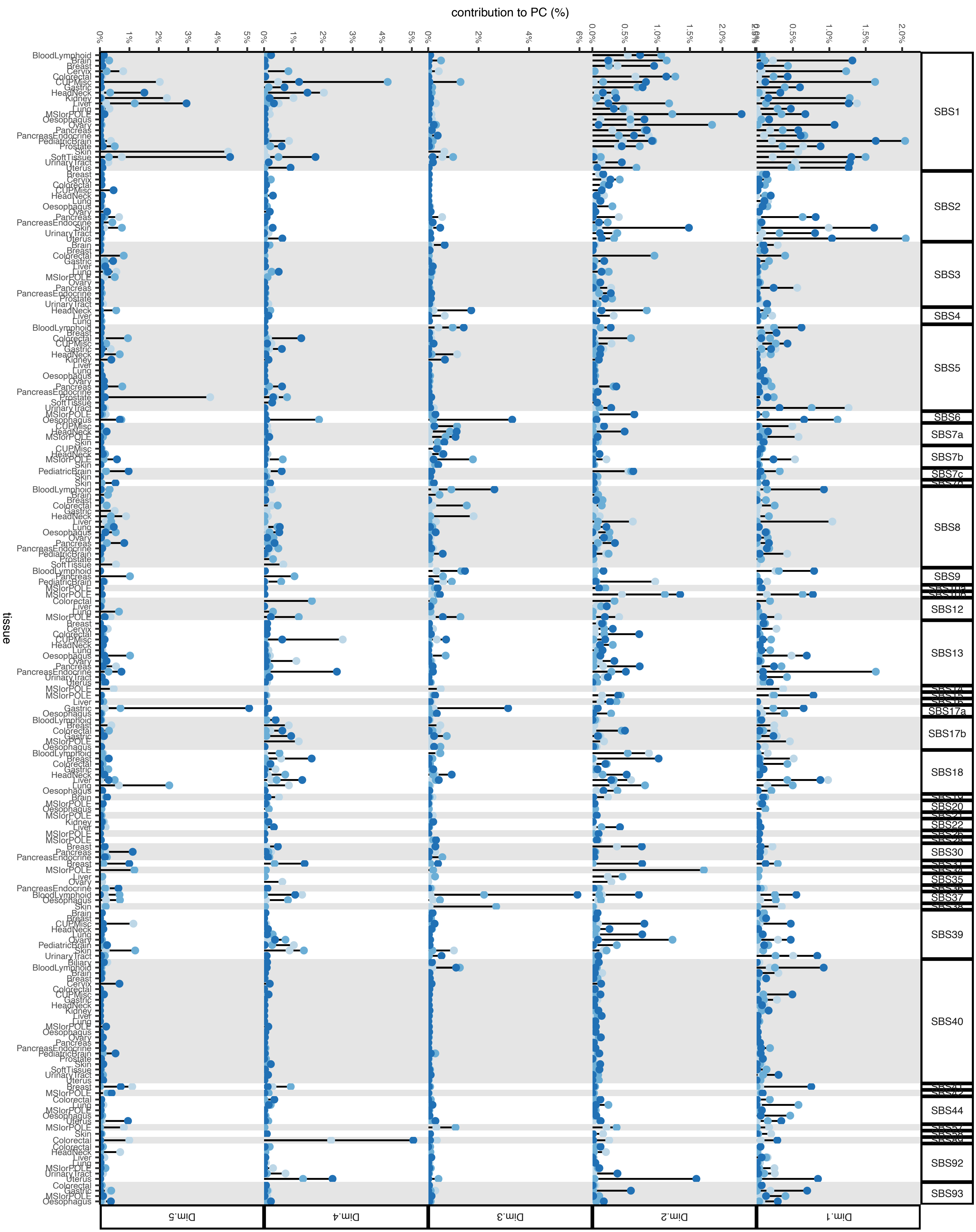

### Supplementary Figure 4

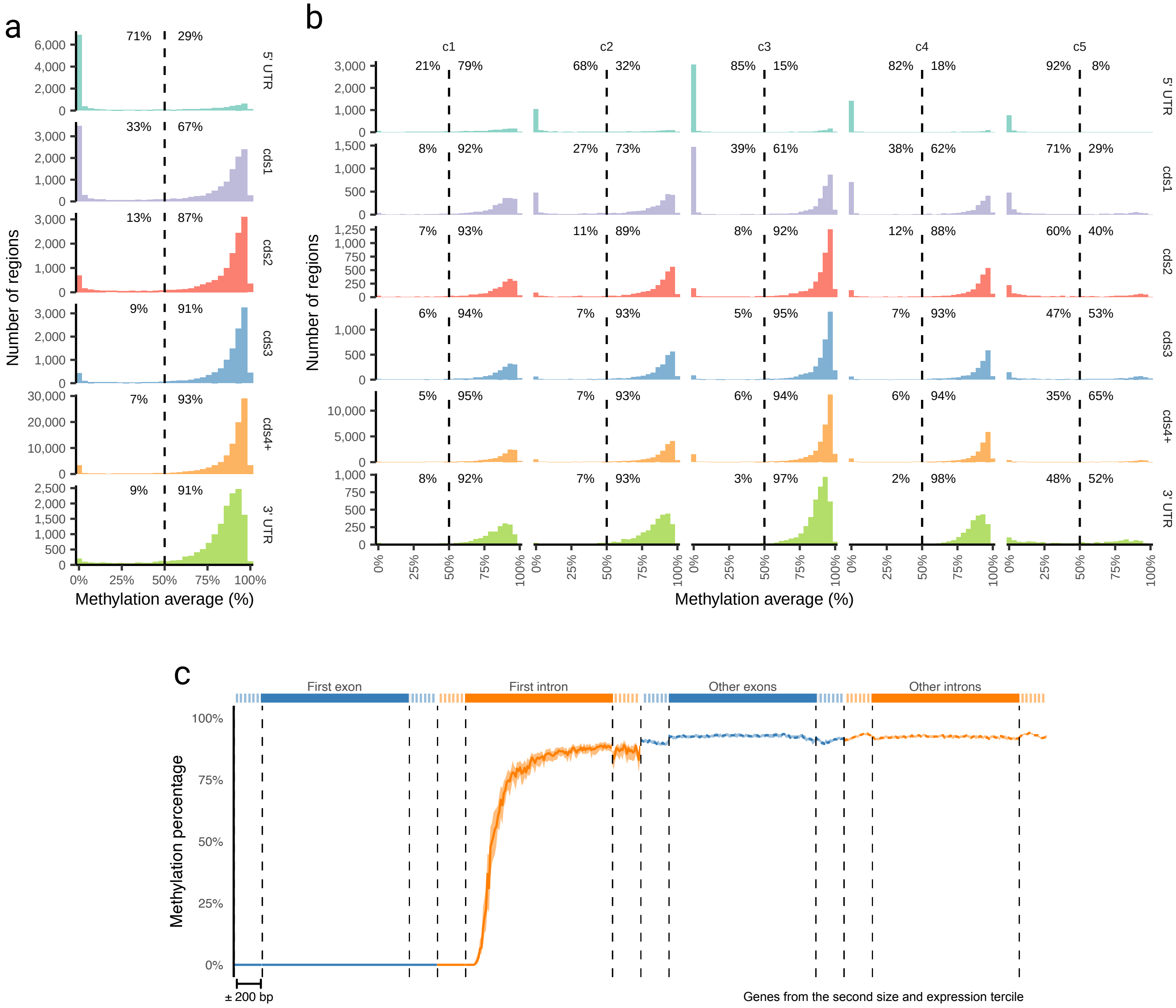

### Supplementary Figure 5

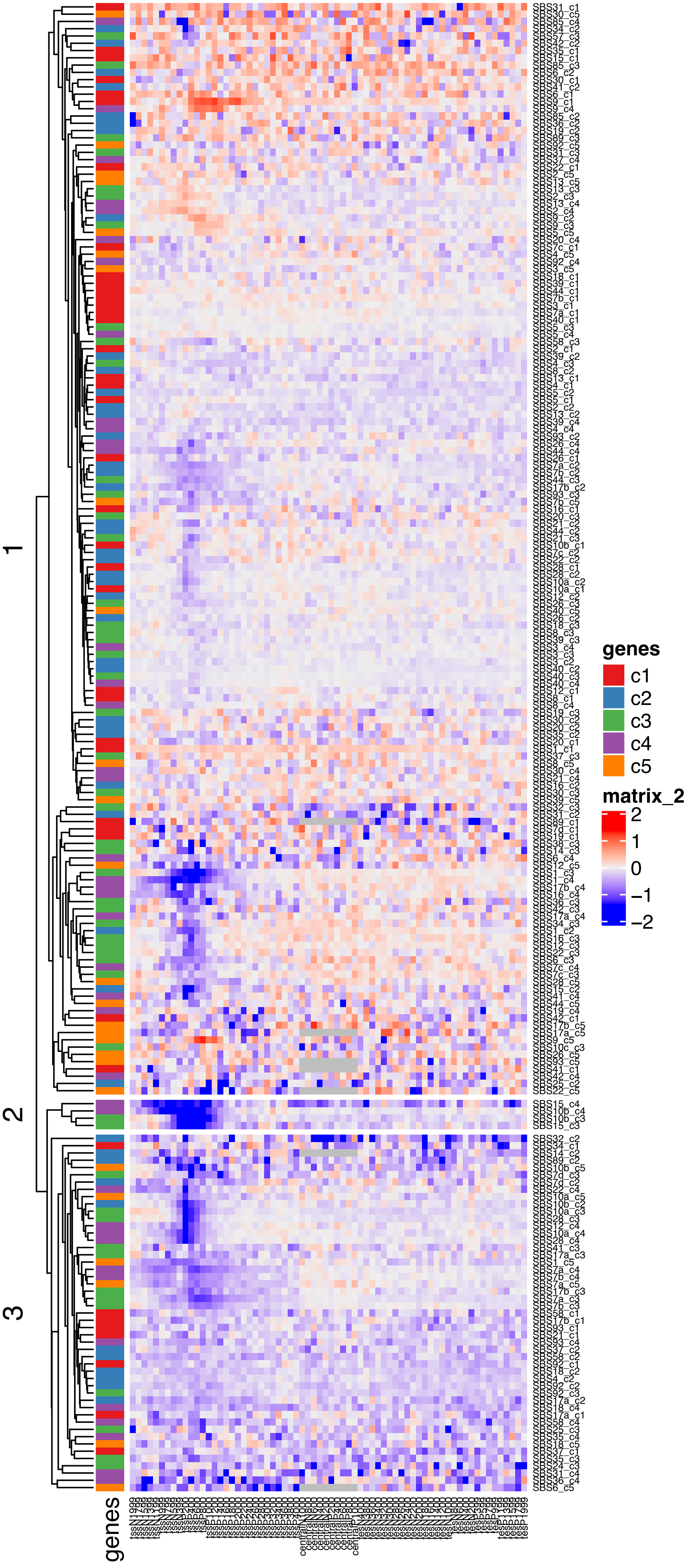

### Supplementary Figure 6

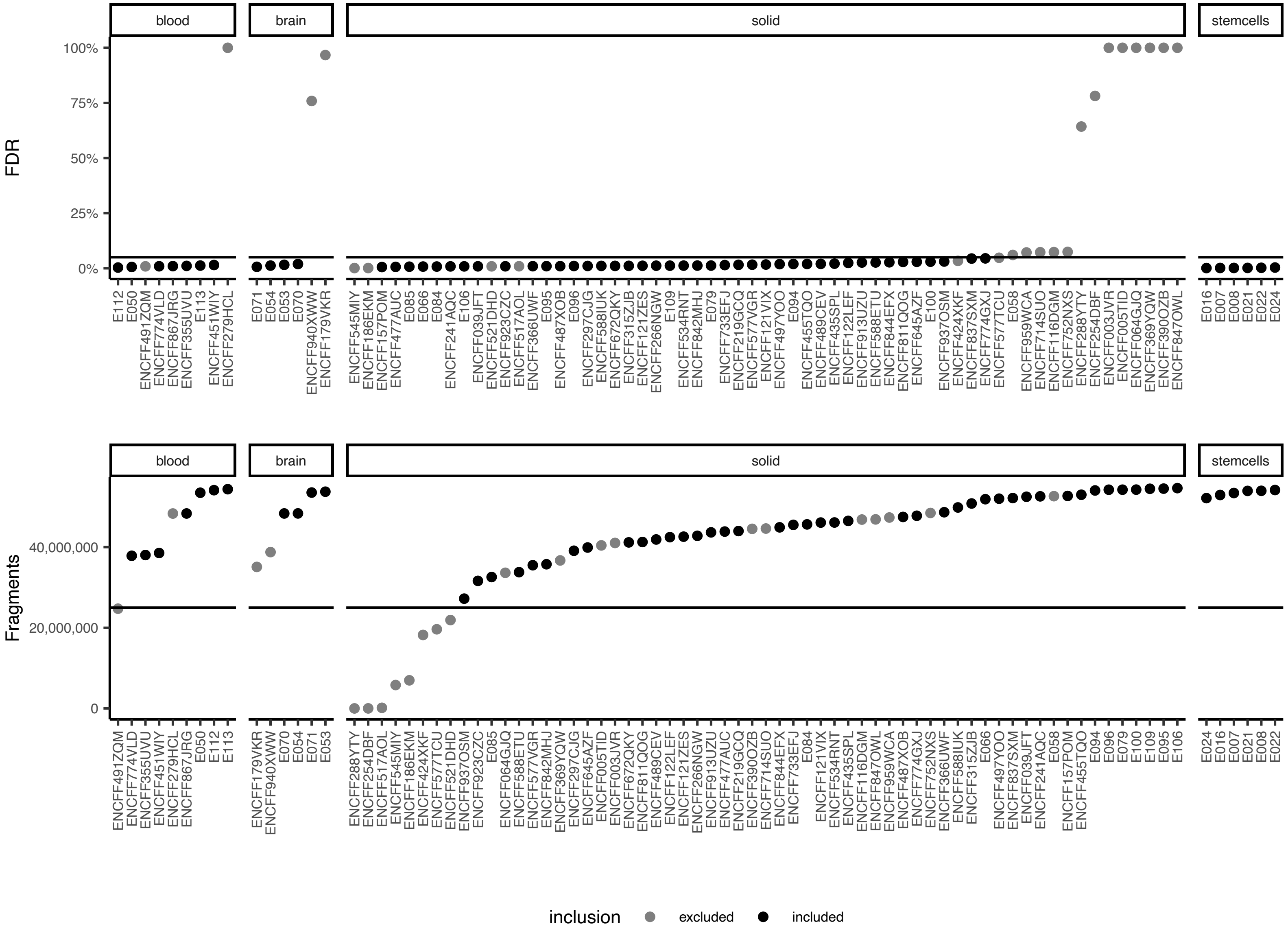

### Supplementary Figure 7

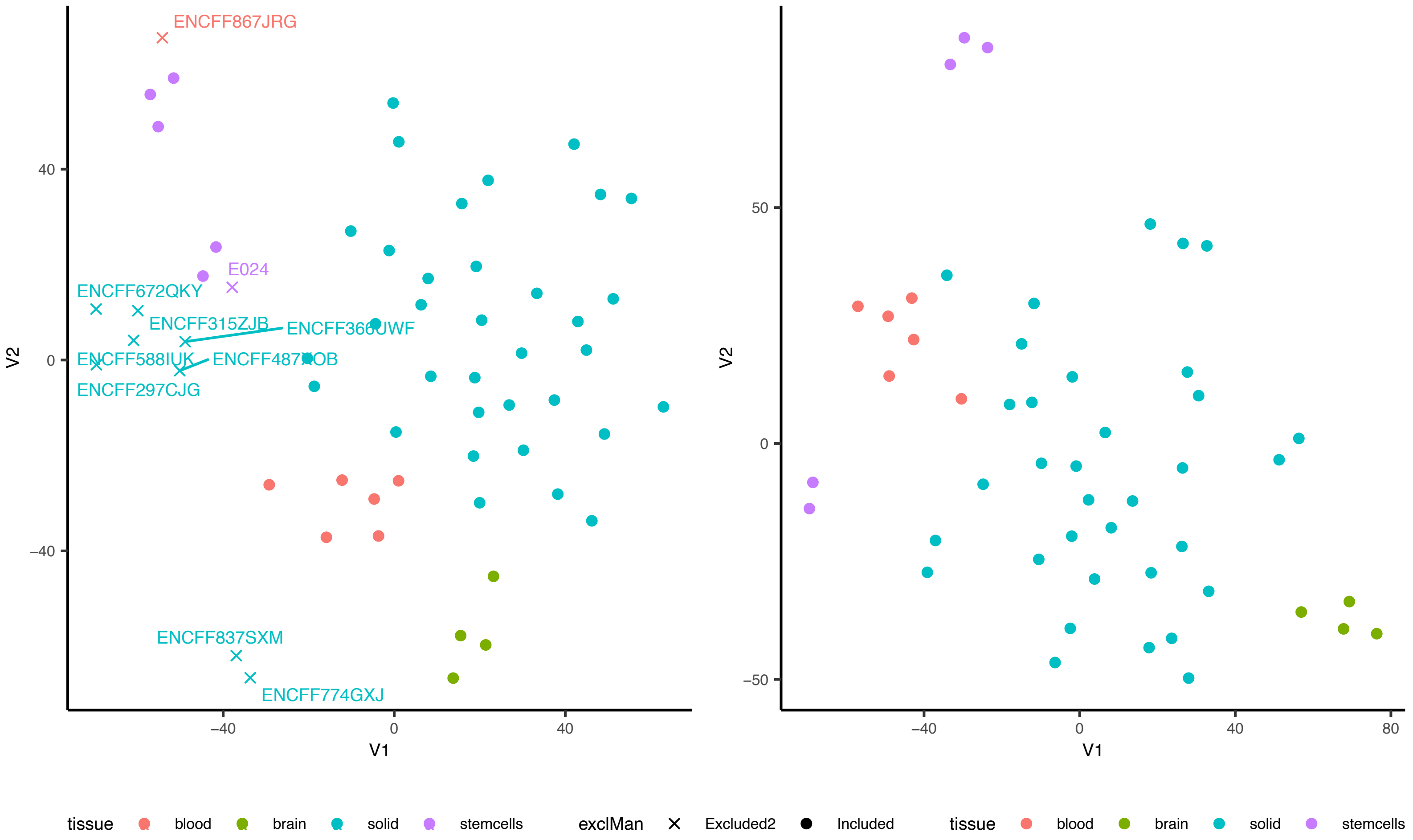
